# Temporal, genome-scale analysis of *Myxococcus xanthus* developmental fate in a mixed population

**DOI:** 10.64898/2026.08.28.747804

**Authors:** Sheenu Mittal, Saikat Mandal, Mark Farrugia, Sean Crosson, Aretha Fiebig, Lee Kroos

## Abstract

*Myxococcus xanthus* bacteria form aggregates when starved on solid surfaces and some cells differentiate into spores. Studies of mutants in monoculture have advanced knowledge of this multi-cellular developmental process, but our understanding of the genetic determinants is incomplete. To assess gene function genomewide, we generated a pool of barcoded transposon insertion mutants, subjected it to starvation, and separated developmental samples into non-aggregated cells, aggregated cells, and spores. We also subjected our pool to chemically-induced unicellular sporulation. Evaluation of changes in the abundance of mutants in samples allowed identification of 200 genes in which insertions reproducibly caused distinct patterns of depletion and/or accumulation over time. Many of these genes have well-established roles in development, validating our approach, while many others have not previously been associated with development. Genes involved in type IV pili (T4P)-dependent motility were more important than gliding motility genes for aggregation and sporulation in the mixed population. Although exopolysaccharide (EPS) synthesis genes are required for aggregation in monoculture, most were dispensable for aggregation in our pool, consistent with EPS sharing between cells, yet these genes were required cell-autonomously for efficient sporulation. Genes for positive regulators of EPS synthesis were important for aggregation as well as sporulation, suggesting functions beyond EPS production. Insertions in several novel genes impaired both starvation- and chemically-induced sporulation. Many genes increased the efficiency of starvation-induced sporulation. Some of these mutants, which we call “developmental winners”, are novel cheaters. Our results demonstrate the power of using the newly-created mutant library to elucidate *M. xanthus* biology.

**IMPORTANCE:** How cells coordinate their activities to build multicellular structures with differentiated cell types is a fundamental question in developmental biology. Starvation triggers thousands of *M. xanthus* cells to move coordinately and build mounds in which some cells differentiate into spores, while other cells lyse or persist as rods. We tracked a barcoded transposon mutant library through development with separation of sub-populations based on aggregation and sporulation fates. We discovered gene sets with distinct abundance profiles over time and across sub-populations. Sets contained both known and uncharacterized genes. For known genes, comparison of our results in a developmentally-competent mixture of mutants with published results for mutants in monoculture distinguished social from cell-autonomous functions. The novel genes provide numerous avenues toward deeper understanding of cellular interactions and differentiation. The mutant library offers a platform for further studies aimed at dissecting *M. xanthus* behaviors functionally, ecologically, and evolutionarily.

## INTRODUCTION

The soil-dwelling bacterium *Myxococcus xanthus* has long served as a model for investigating mechanisms that govern bacterial multicellularity and differentiation (1). Under nutrient-limiting conditions, *M. xanthus* initiates a developmental program that leads to the formation of multicellular fruiting bodies, which are specialized structures where motile, rod-shaped cells differentiate into stress-resistant myxospores. This developmental program is fundamentally social: cells cooperate through cell-cell signaling and shared extracellular products. Only a minority of the starving population survives to become spores, and the fate of any individual cell is influenced by the genotypes surrounding it.

Genetic studies in *M. xanthus* over the past several decades have uncovered numerous transcriptional regulators and cell-cell signaling pathways that orchestrate the temporal and spatial transitions required for fruiting body development (1, 2). However, our understanding of *M. xanthus* development is incomplete, and significant opportunities remain to discover new genes and molecular processes that govern multicellular aggregation and spore differentiation. Notably, select chemical treatments can induce rapid, unicellular *M. xanthus* sporulation (3), which can proceed *via* mechanisms differing from the sporulation pathway that operates during starvation-induced fruiting body formation (4, 5). Studies of alternative sporulation routes provide an opportunity to expand understanding of spore formation and reveal novel regulators and effectors of important developmental processes.

To comprehensively identify genes that contribute to *M. xanthus* development, we conducted a genome-scale, time-resolved analysis using a randomly barcoded transposon mutant library. This diverse collection of mutants develops as a pool, so each mutant proceeds through development as a rare genotype within a population of largely wild-type-like cells, allowing us to assess how loss of a given gene affects an individual cell competing within a community. By physically separating non-aggregated cells, aggregated cells, and mature spores across time, we resolved where in development each mutant was depleted or enriched, distinguishing effects on attachment and/or survival, aggregation, and sporulation. In parallel, we evaluated our mutant pool after glycerol-induced sporulation.

This approach revealed expected fitness profiles for previously characterized developmental genes, validating our approach, and also uncovered numerous new genes that impact fruiting body development and/or glycerol-induced sporulation. The fate-resolved, mixed-population design also distinguished cell-autonomous requirements from functions shared among cells, and identified genes whose loss compromises attachment and/or survival during development. It further revealed developmental winners, mutants that sporulate more efficiently than their neighbors, linking gene function to cheating and phase variation in some cases. Together, these results provide novel insights into the genetic underpinnings of multicellular development and alternative spore differentiation pathways in bacteria, illuminate how rare individual genotypes fare within social groups, and define a valuable framework for future functional dissection and experimental evolution of developmental processes in *M. xanthus*.

## RESULTS AND DISCUSSION

### A BarSeq approach to define genetic requirements for *M. xanthus* development

To enable genetic analysis of *M. xanthus* development, we generated a pool of barcoded Tn-Himar insertion mutants for random barcode transposon insertion site sequencing (RB-TnSeq or BarSeq) (6). Mapping the insertion sites in the pool identified 123,216 reliably mapped barcodes at 61,040 distinct sites in the genome. This *M. xanthus* Tn-Himar pool contains a median of 6 mutant strains per protein, and at least one insertion in the central region of 81% of protein-coding genes.

We used the barcoded Tn-Himar mutant pool to monitor changes in mutant representation during submerged-culture development under starvation (Fig. 1). The pool was grown to log phase, transferred from complex medium into MC7 buffer, incubated in 6-well plates to initiate development, and harvested across a developmental time series. The vast majority of cells formed a biofilm attached to the well bottom. At the time of harvest, the culture supernatant was carefully replaced with fresh MC7 buffer, so the few (if any) unattached cells were discarded and the attached biofilm cells were kept for analysis. In the first experiment (time course 1), we analyzed the unfractionated population at early time points (12 to 36 h), capturing non-aggregated (non-agg) and aggregated (agg) cells together. In two additional experiments (time courses 2 and 3), we extended sampling into later developmental stages (up to 72 h) and separated developmental fates: non-agg cells were recovered from the supernatant after a low-speed spin and agg cells from the pellet. From separate pellet samples, spores were purified by breaking non-spore cells, digesting the released DNA with DNase I, inactivating the enzyme, and then breaking spores with glass beads. In parallel, we induced sporulation by adding glycerol and processed the resulting spores in the same way. Together, these experiments were designed to identify genes required for attachment and/or survival through early development, for aggregation versus remaining non-aggregated, and for spore formation across distinct sporulation contexts (Fig. 1A and 1B).

**FIG 1.**
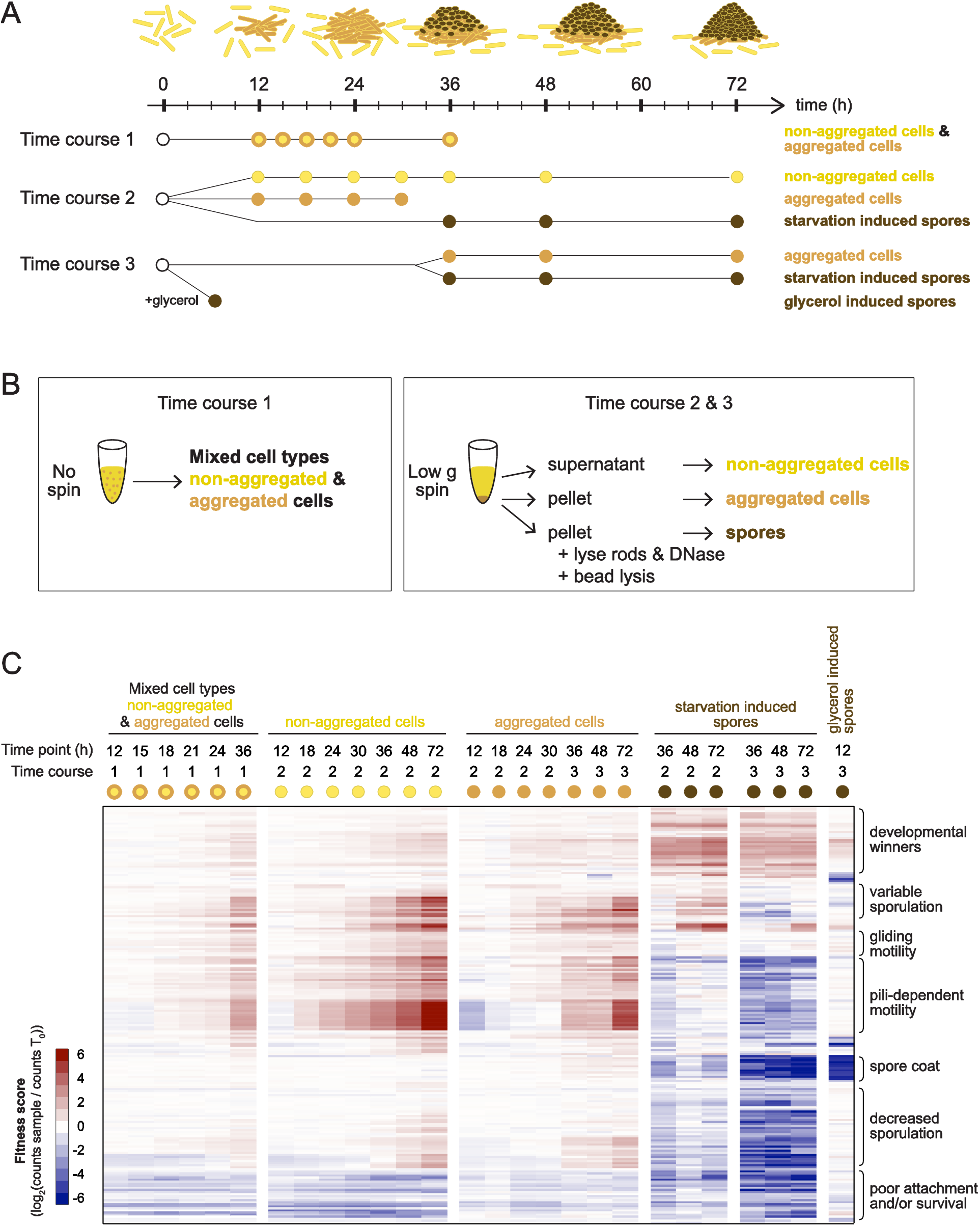
Barcoded TnSeq reveals genes impacting *M. xanthus* development. (A) Experimental overview. (B) Approaches to fractionate cell types. (C) Hierarchical cluster of fitness scores of 200 genes with an absolute average t-score >4 in at least one biological replicate sample and an absolute average fitness score >1 in at least one complete sample. Brackets indicate groups with distinct fitness profiles.

For each time point and fraction, we amplified, sequenced, and counted barcodes, then used these counts to compute gene-level fitness scores and t-scores following the approach of Wetmore and colleagues (6). Fitness scores express the change in relative abundance of barcodes associated with transposon insertions in a given gene, on a log2 scale, relative to the input (0 h) sample, while t-scores quantify how consistently mutants carrying independent insertions in that gene behave. Given that these assays were performed under starvation or glycerol spore-inducing conditions rather than during growth, barcode dynamics should largely reflect developmental fate partitioning rather than population expansion through cell division. This experimental context impacts how we interpret changes in barcode (i.e. mutant) relative abundance. Moreover, a majority of starving cells die during development accompanied by chromosomal DNA degradation (7-10), though whether this death is a programmed developmental fate remains unclear. Nevertheless, barcodes showing disproportionate loss early in development should mark genes important for attachment and/or survival.

To attribute developmental phenotypes to genes robustly, we prioritized genes supported by multiple independent Tn-Himar insertions with sufficient read depth and reproducible effects across replicates, applying standard thresholds to call significant mutant depletions or enrichments. Genes with an absolute average t-score >4 in any biological replicate sample from time courses 1, 2, or 3, and also an absolute average fitness score >1 in any complete sample (i.e., 2 biological replicates, each with 3 technical replicates), were then subjected to hierarchical clustering, which resolved fraction-specific fitness patterns across developmental time points. Distinct mutant groupings were enriched or depleted in non-agg cells, agg cells, or purified spores, reflecting altered developmental fate partitioning (Fig. 1C, Table S1).

### T4P-dependent motility is more important than gliding motility for aggregation and sporulation

Starving *M. xanthus* coordinate their movements to build mound-shaped aggregates during fruiting body development. Two motility systems control cell movement (1) (Fig. 2A). Social or “S” motility involves movement of cells in groups *via* cycles of extension, adhesion to neighboring cells or the substratum, and retraction of T4P. Adventurous or “A” motility involves gliding of individual cells *via* transient interactions of bacterial focal adhesion complexes (bFACs) with the substratum. The bFACs span the cell envelope and convert the proton-motive force across the inner membrane into mechanical force for movement, likely by pushing against cytoplasmic MreB filaments and periplasmic peptidoglycan (PG). Both motility systems assemble at the leading pole to move the cell in one direction and, periodically, switch to the opposite pole, reversing the direction of movement. Regulation of motility reversal involves the Frz chemosensory system feeding into a multi-protein polarity module. Early work suggested that S-motility mutants aggregate abnormally, while most A-motility mutants aggregate normally (11). Later studies defined the genes and molecular mechanisms underlying each motility type (1), but only recently has the development of a motility mutant been examined in the context of excess cells with normal motility (12). Our approach extends this comparison to many motility mutants at once within the Tn-Himar pool, in which the vast majority of insertion mutants have no significant developmental defect.

**FIG 2.**
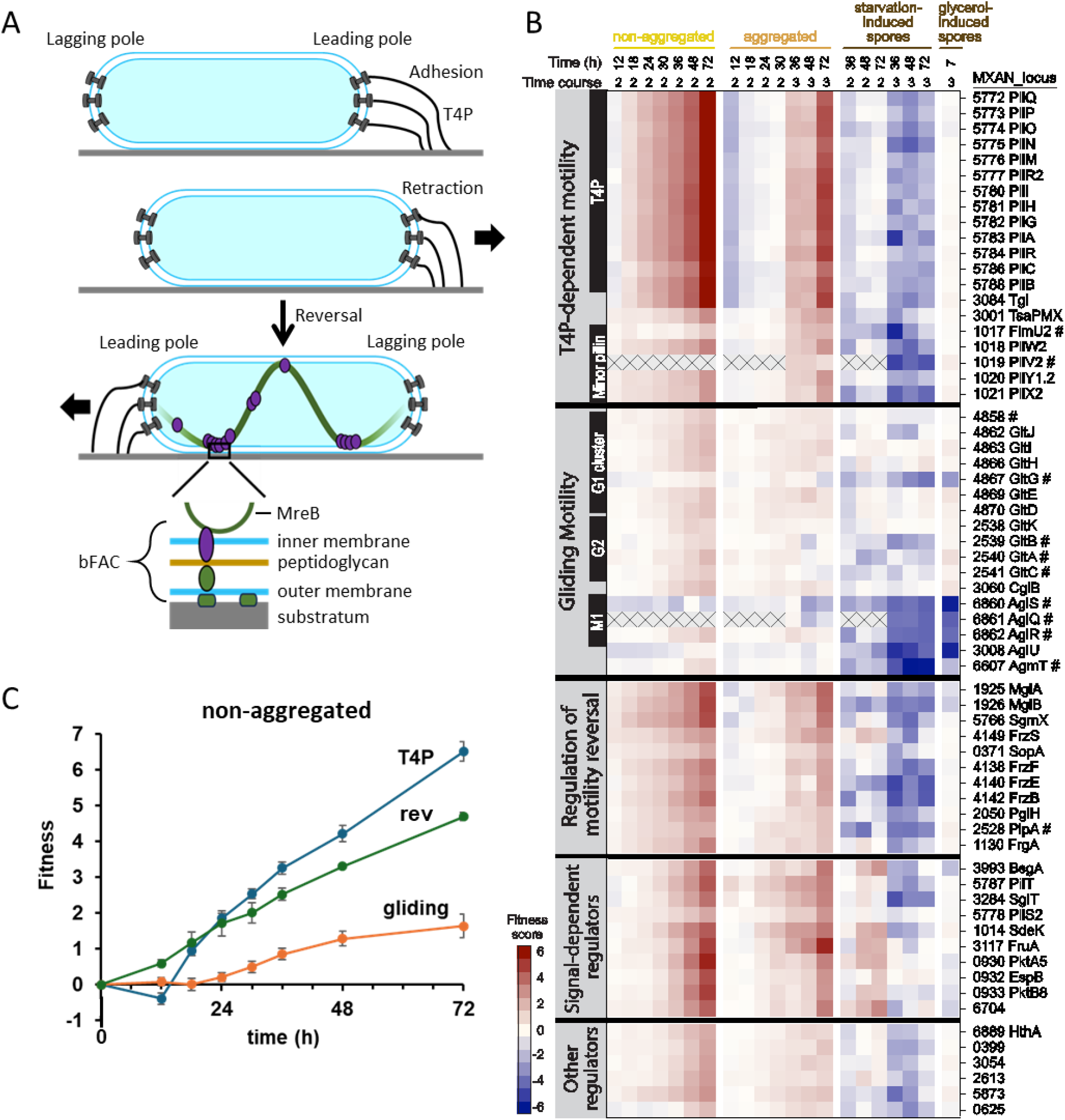
Motility and signal-dependent regulators are critical determinants of starvation-induced development. (A) Type IV pili (T4P) extend from the leading cell pole, adhere to the substratum, and retract. Cells reverse movement by switching T4P to the opposite end coordinately with bacterial focal adhesion complexes (bFACs) shown only in the bottom cell. Inset shows a bFAC spanning from cytoplasmic MreB filaments across the cell envelope to the substratum to power gliding motility. (B) Heat map showing fitness scores of genes for motility, signal-dependent regulators, and other genes with a similar fitness profile. #; gene not among the 200 genes shown in Figure 1C and Table S1. Gray boxes with X indicate samples from time courses with insufficient counts at time 0 to assign scores. (C) Average fitness scores for genes involved in T4P-dependent motility (13 *pil* genes and *tgl*), regulation of motility reversal (rev; *mglA*, *mglB* and *sgmX*), and gliding motility (most *glt* genes except *gltG*, and *cglB*) in non-aggregated samples over time. Error bars show one standard deviation.

#### T4P-dependent motility

Most proteins for T4P-dependent motility are encoded at the 17-gene *pil* locus (MXAN_5772-5788). Tracking of *pilC* cells mixed with a 10,000-fold excess of wild type previously showed poor rescue of *pilC* aggregation and sporulation (12), so we expected *pil* insertion strains to shift distribution across developmental fates in experiments with our Tn-Himar pool. Strains with insertions in *pilC* (MXAN_5786) and 12 other *pil* genes shared strikingly similar fitness patterns (Fig. 1 and 2B; Table S1). At early time points (12, 18 h) these strains were underrepresented in agg samples (fitness scores approximately −2 at 12 h; Fig. 2B and S1), indicating an aggregation defect, and revealing that T4P contribute to aggregation even before visible mound formation. These mutants progressively accumulated in non-agg samples, reaching nearly 100-fold overrepresentation by 72 h (fitness scores +6 to +7), underscoring the importance of T4P-dependent motility for aggregation. Their negative scores in starvation-induced spore samples align with the poor sporulation of a *pilC* mutant co-developed with excess wild type (12), and of a *pilA* mutant alone in submerged culture (13). Unexpectedly, *pil* mutants showed positive fitness scores in agg samples at 36 h and later (Fig. 2B and S1). The scores were lower than in non-agg samples at those time points, a pattern observed for most mutants (Fig. 1), suggesting that our protocol did not completely remove non-agg cells from agg samples. We note that genes at this locus known to regulate T4P retraction (*pilT*) or assembly (*pilS2*) exhibit distinct fitness patterns, described below.

Tgl is an outer membrane (OM) lipoprotein required for T4P formation (14). Tgl can be transferred from *tgl^+^* cells to *tgl^-^* cells (15) *via* OM exchange (OME) (16), allowing transient “stimulatable” S-motility of *tgl^-^* cells during growth (11). However, insertions in *tgl* (MXAN_3084) resulted in a similar fitness profile as the 13 *pil* genes (Fig. 2B; Table S1), indicating Tgl transfer did not rescue the development of *tgl* mutants in our mutant pool. We conclude that during development Tgl functions cell-autonomously to allow T4P-dependent motility.

T4P-dependent motility also depends on TsaPMX and minor pilins (17, 18). During growth, TsaPMX localizes to the cell poles and aids T4P surface assembly (17), while minor pilins prime pilus assembly and localize to the tip, where they are thought to mediate adhesion (19). Either cluster 1 or cluster 3 minor pilins are sufficient for T4P-dependent motility in growing cells, but cluster 2 minor pilins fail to accumulate detectably in cells during growth (19). All five cluster 2 genes, however, are transcriptionally up-regulated during development (20-23); no developmental role for TsaPMX or minor pilins had been reported. We observed that insertions in *tsaPMX* (MXAN_3001) and cluster 2 genes *pilW2*, *pilY1.2*, and *pilX2* (MXAN_1018, 1020, 1021) showed fitness patterns similar to, but weaker than, the 13 *pil* genes and *tgl*. The other two cluster 2 genes, *fimU2* (MXAN_1017) and *pilV2* (MXAN_1019), behaved similarly (Fig. 2B; Table S2) but fell below our statistical cutoffs for Figure 1 and Table S1. We conclude that cluster 2 minor pilins and TsaPMX are important for T4P function during development, revealing specialized developmental roles for distinct minor pilins.

#### Gliding motility

Most bFAC gliding-machinery proteins are encoded at three loci (24): the G1 (MXAN_4862-4870) and G2 (MXAN_2538-2541) clusters encode Glt proteins that associate with the AglS/Q/R motor (MXAN_6860-6862) encoded at the M1 locus. The inner-membrane AglS/Q/R motor harnesses the proton-motive force to power gliding with the Glt proteins and spore-coat formation with the Nfs proteins (25). Nfs proteins are required for coat deposition during both starvation- and glycerol-induced sporulation (5, 26). Given these previous studies, we expected most insertions in genes at the three loci to have little or no effect on fitness for developmental aggregation (11) and insertions in the M1 cluster to strongly reduce sporulation (25).

Unexpectedly, insertions in most G1 and G2 *glt* genes, and in *cglB* (MXAN_3060), a gliding-machinery gene at a separate locus, gave fitness patterns resembling those of T4P genes but with dampened magnitude (Fig. 2B-C and S1). Additionally, MXAN_4858, an uncharacterized gene near the 3′ end of the G1 cluster, behaved similarly, though an insertion in this gene was previously reported to have no overt motility effect (27). MXAN_4858 contains a TIGR04552 domain that is broadly conserved in the Myxococcales, and may function in concert with other gliding-motility genes to modestly promote development. Collectively, these results are consistent with A-motility contributing to aggregation, but less than S-motility (11). Our data also permit quantitative comparison of motility loss within a mixed population. By the end of the time course, the 13 *pil* genes and *tgl* were overrepresented ∼100-fold (2^6.5^) in non-agg cells, whereas *cglB* and most *glt* mutants (except *gltG*; see below) were overrepresented only ∼3-fold (2^1.6^) (Fig. 2C). Similarly, disruption of the T4P machinery was more detrimental to spore formation than disruption of the gliding machinery (Fig. 2B).

CglB, CglC (GltK), and CglE (GltH) are OM proteins that, like Tgl, undergo OME (15, 16). CglB/C/E transfer confers stimulatable A-motility during growth (28). *cglB/C/E* insertion mutants resembled most *glt* mutants (Fig. 2B), indicating that CglB/C/E OME did not rescue developmental gliding and that, like Tgl, these proteins act cell-autonomously. This is consistent with the normal monoculture development of an OME-deficient *traA* mutant (29).

Insertions in the M1 genes *aglS/Q/R* were strongly depleted from glycerol-induced spore samples, as expected (25), and also from starvation-induced spore samples (Fig. 2B; Table S2). *aglS* mutants were additionally underrepresented in most non-agg and agg samples, while *aglQ* was too sparsely represented in time course 2 to score. AlphaFold modeling predicts that AglS and AglQ form a heterodimer plugging the pentameric AglR proton channel (30). We propose that failure to plug this channel impairs *aglS* and *aglQ* mutant survival, explaining their low overall fitness (Fig. 2B). In contrast, *aglR* mutants were modestly enriched in non-agg and agg samples, comparable to most *glt* mutants.

Loss of the proton channel is expected to be less deleterious than a leaky proton channel and thus we infer that the fitness phenotypes of the *aglR* mutants in non-agg samples likely result from modest aggregation defects due to loss of gliding-motility power.

Surprisingly, *gltG* (MXAN_4867) mutants resembled *aglS/Q/R* mutants more than other *glt* or *cglB* mutants: they showed almost no effect in non-agg and agg samples but were underrepresented in both starvation- and glycerol-induced spore samples (Fig. 2B; Table S2). GltG interacts directly with AglR (24, 30), as does its Nfs-system homolog NfsG (MXAN_3377) (25). The spore depletion phenotype therefore suggests a role for GltG in coat deposition, while the lack of an aggregation phenotype suggests that NfsG substitutes for GltG during aggregation (Fig. 2B). Hence, a *gltG nfsG* double mutant may exhibit more severe aggregation and sporulation defects than either single mutant.

Two other genes that contribute to gliding motility, *aglU* (MXAN_3008) and *agmT* (MXAN_6607), had fitness score profiles that indicate roles in both sporulation pathways. *aglU* is predicted to encode an OM lipoprotein that may mediate assembly of protein complexes (31). Similar to *aglS*, *aglU* insertion strains were modestly underrepresented in non-agg and agg samples (Fig. 2B; Table S1) suggesting a role in attachment and/or survival. *aglU* mutants were strongly depleted in spore samples, similar to *aglS/Q/R* mutants, implying a role in coat formation and consistent with the immature starvation-induced spores reported for *aglU* mutants (31). The fitness profile of *agmT* was similar to *aglR* (Fig. 2B; Table S2). AgmT is a lytic transglycosylase that likely modifies PG and connects bFACs to PG, enabling force generated by AglR/Q/S motors to push against PG and power gliding motility (32). Although a defined *agmT* mutant in monoculture showed no sporulation defect (33), *agmT* mutants in our pool were defective in both sporulation pathways, expanding the functional role of this PG-modifying enzyme.

#### Regulation of motility reversal

MglA/B are key components of the polarity module that controls reversal of cell movement in response to the Frz chemosensory system (1). MglA is a GTPase that when bound by GTP defines the leading cell pole, and MglB is its cognate GTPase-activating protein (34, 35). Since this module coordinately switches both motility systems (Fig. 2A), and *pil* and *tgl* insertions affected development more strongly than *glt* and *cglB* insertions (Fig. 2B and 2C), we expected insertions in *mglA/B* and crucial downstream T4P effectors to strongly affect developmental fitness profiles. SgmX, FrzS, and SopA are downstream effectors of the polarity module, with differential importance for T4P-dependent motility (36). MglA interacts directly with SgmX (37, 38), which in turn stimulates PilB localization at the leading pole for extension of T4P (38). FrzS and SopA stimulate the MglA/SgmX pathway to differing degrees (36).

As expected, insertions in *mglA/B* (MXAN_1925/1926) and *sgmX* (MXAN_5766) produced fitness profiles resembling the 13 *pil* genes and *tgl* (Fig. 2B and 2C; Table S1), supporting key roles for MglA/B and SgmX in aggregation and sporulation within the mixed population. The congruent *mglA* and *mglB* mutant fitness patterns likely reflect polar effects of *mglB* insertions on *mglA* (39). The developmental phenotype of an in-frame *sgmX* deletion has not been reported; although *sgmX* appears to be the first gene of an operon, insertions in downstream genes (MXAN_5765-5761) did not exhibit developmental defects (Table S2), indicating that the phenotype is specific to *sgmX* rather than polar. Insertions in *frzS* (MXAN_4149) and *sopA* (MXAN_0371) caused progressively weaker accumulation in non-agg samples than insertions in *mglA/B* and *sgmX* (Fig. 2B; Table S1). This implies a hierarchy of effects on aggregation: MglA/B and SgmX > FrzS > SopA. This gradient matches the quantitatively defined effects of these proteins on T4P-dependent motility (36), showing that our genome-scale fitness measurements sensitively distinguish phenotypes in the mixed population. The same ranking held for starvation-induced sporulation, where *frzS* and *sopA* insertions again had less effect than *mglA/B* and *sgmX* (Fig. 2B). Notably, *frzS* mutants (like *pilT* mutants) were overrepresented in starvation-induced spore samples in time course 2 but underrepresented in time course 3 (Fig. 2B), a pattern shared by several genes implicated in signal-dependent gene regulation (see below).

Upstream of the polarity module, the Frz chemosensory system regulates motility reversal (1). Early work identified *frz* mutants that form aggregates with a macroscopic “frizzy” filamentous appearance (40) due to infrequent reversal of motility (41). Mutants with insertions in *frzF/E/B* (MXAN_4138/4140/4142) accumulated in non-agg samples slightly less than mutants with insertions in *mglA/B* and *sgmX* (Fig. 2B; Table S1), and were similarly depleted from starvation-induced spore samples. Our results are consistent with previous studies showing abnormal frizzy aggregation of *frzF/E/B* in-frame deletion mutants (42) and severely impaired sporulation of *frzF* insertion mutants (43). Since FrzF/E/B appear less important for aggregation than MglA/B and SgmX yet at least as important for starvation-induced sporulation (Fig. 2B), we postulate that they act primarily at a late developmental step, such as the intra-aggregate cell movements that enhance spore formation (44). Although the developmental signals to which the Frz chemosensory system responds remain elusive, the lipid phosphatidylethanolamine is a Frz-dependent chemoattractant (45), and other lipids are intercellular signals that may act primarily at short range during development (46-48). Insertions in *pglH* (MXAN_2050), *plpA* (MXAN_2528), and *frgA* (MXAN_1130) produced patterns similar to *frzF/E/B* (Fig. 2B; Tables S1 and S2). Some evidence exists that these genes contribute to regulation of motility reversal (see SI), though further work is needed to clarify their role in this regulation.

### Known signal-dependent regulators and uncharacterized genes reduce aggregation and sporulation

A signaling and gene regulatory network controls the expression of well over 1000 genes to govern *M. xanthus* aggregation and sporulation (1, 2). BsgA is a Lon-family ATP-dependent protease implicated in extracellular signal production (49, 50), but the "B-signal" it is presumed to generate has not been identified, and attempts to demonstrate extracellular signaling by co-developing *bsgA* mutants with wild type have given inconsistent results (50-52). Nevertheless, these studies agree that *bsgA* mutants exhibit defects in early developmental gene expression, aggregation, and sporulation. In our experiments, *bsgA* (MXAN_3993) insertions produced a fitness pattern resembling those of the T4P retraction gene *pilT* (MXAN_5787) and the reversal-regulating gene *frzS*: enrichment in non-agg and agg samples indicating defective aggregation, and variable abundance in spore samples (Fig. 2B; Table S1; SI). We speculate that insertions in *bsgA*, *pilT*, *frzS*, and eight other genes (see below) cause starvation-induced sporulation to vary with conditions that differed between our two experiments. Such sensitivity to assay conditions could also explain the inconsistent results of co-developing *bsgA* mutants with wild type (50-52). Notably, our data reveal a defect in glycerol-induced sporulation of *bsgA* mutants (Fig. 2B), a novel observation meriting further investigation.

Eight other genes had fitness profiles similar to *bsgA*, *pilT*, and *frzS*. Seven of these, *sglT* (MXAN_3284), *pilS2* (MXAN_5778), *sdeK* (MXAN_1014), *fruA* (MXAN_3117), and the *pktA5*-*espB*-*pktB8* cluster (MXAN_0930, 0932, 0933), encode known signal-dependent regulators of development (Fig. 2B; Tables S1 and S2; see SI for expanded discussion of these genes). The eighth, MXAN_6704, encodes a predicted Gcn5-related N-acetyltransferase (GNAT) with no known substrate. An MXAN_6704 mutant produces fewer lipid bodies than normal during development, indicating a defect in converting membrane phospholipids to the triacylglycerides stored in lipid bodies and consumed during spore maturation (53). This mutant is also defective in starvation-induced aggregation and sporulation. Glycerol-induced sporulation was not previously tested, but our experiments show that insertions in MXAN_6704 cause a defect (Fig. 2B; Table S1).

Together, insertions in these genes implicated in T4P dynamics and signal-dependent regulation reduced aggregation and yielded variable sporulation outcomes, consistent with developmental functions whose loss renders cells sensitive to assay conditions (and possibly to density or complementation effects) in the mixed population. These interpretations can be tested by co-developing marked mutant and wild-type strains to read out aggregation timing, sporulation efficiency, and developmental gene expression (44, 54).

Insertions in *hthA* (MXAN_6889) and five uncharacterized genes (MXAN_0399, 3054, 2613, 5873, 0625) also produced fitness patterns consistent with reduced aggregation, but unlike the genes above, these mutants were underrepresented in starvation-induced spore samples in both experiments (Fig. 2B; Table S1). HthA is a predicted DNA-binding protein important for aggregation; an *hthA* insertion mutation delayed and reduced sporulation and altered expression of some developmental genes, possibly through a polar effect on the downstream *hthB* gene (55). The five remaining genes encode proteins from large enzyme families with diverse roles (see SI for more details) and have not previously been implicated in development.

### Cell-surface polysaccharides play key roles in development

*M. xanthus* secretes three polysaccharides and embeds a fourth in the surface leaflet of its OM (1, 56) (Fig. 3A), each synthesized and transported across the cell envelope by a separate multi-protein pathway (Fig. 3B). Exopolysaccharide (EPS) remains largely surface-associated, forming a protective layer that also stimulates T4P-dependent motility. Spore coat polysaccharide (SCP) is made only during development and deposited on spores by Nfs proteins driven by the AglS/Q/R motors, as described above. Biosurfactant polysaccharide (BPS) is secreted without remaining cell-associated and primarily affects vegetative colony morphology rather than fruiting body development. Lipopolysaccharide (LPS) occupies the OM surface leaflet and is important for gliding motility and development. Within our Tn-Himar pool, insertions in genes known to be important for EPS, SCP, and LPS production were strongly depleted in starvation-induced spore samples while showing several distinct patterns in other samples, and we additionally identified genes with similar profiles and annotations suggesting roles in polysaccharide production but no prior link to development.

**FIG 3.**
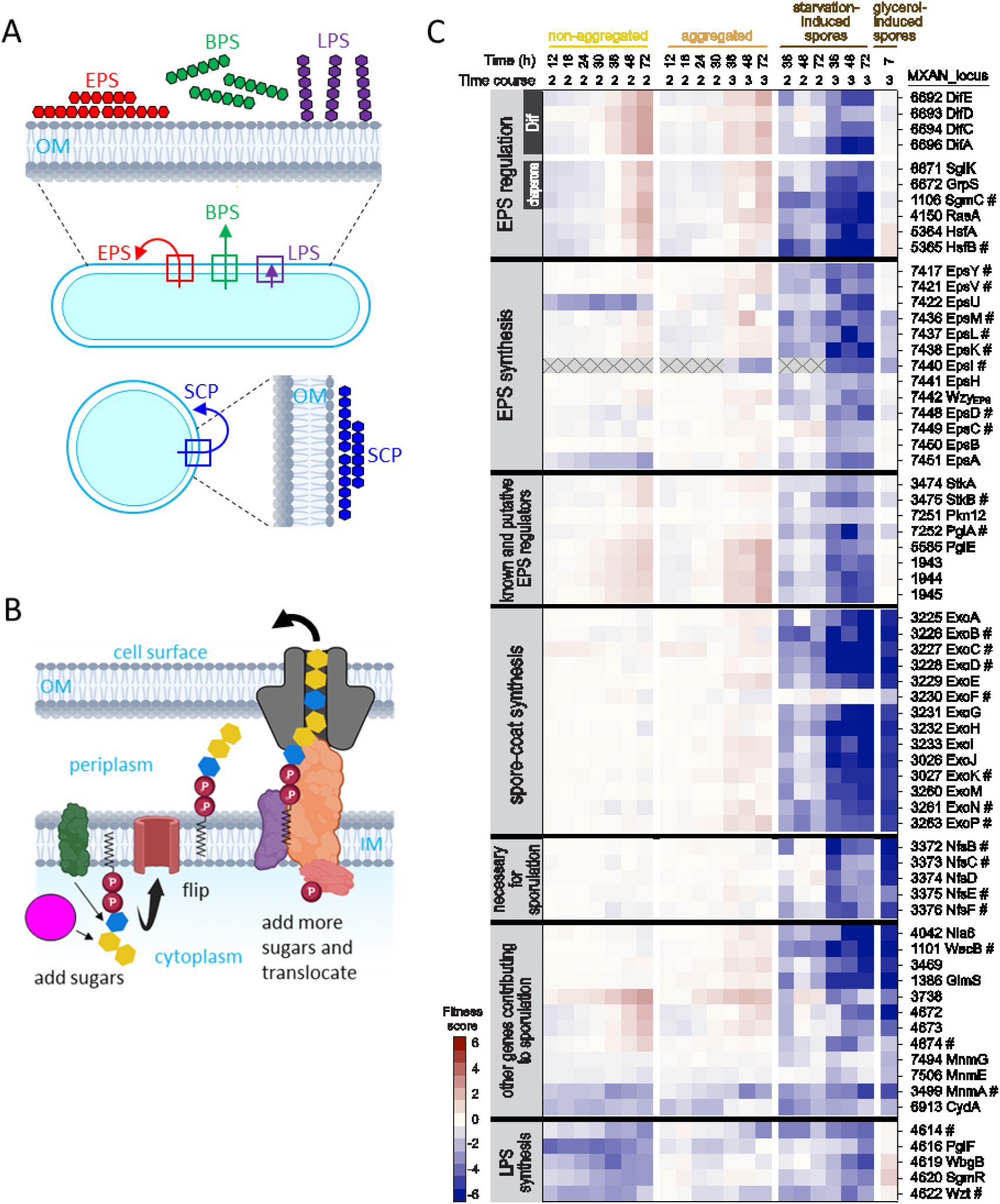
Cell-surface polysaccharides play key roles in development. (A) Rod-shaped cells synthesize exopolysaccharide (EPS), biosurfactant polysaccharide (BPS), and lipopolysaccharide (LPS), and round spores synthesize spore coat polysaccharide (SCP), each *via* a distinct pathway (boxes). Arrows and expanded views depict primary destination of polysaccharide relative to outer membrane (OM). (B) Simplified polysaccharide synthesis pathway. Enzymes add sugars to lipid in the inner membrane (IM), an IM protein flips polysaccharide to the periplasm, IM proteins add more sugars, and an OM protein translocates polysaccharide to the cell surface. (C) Heat map showing fitness scores of genes for EPS regulation and synthesis, SCP synthesis and deposition, other genes important for both types of sporulation, and LPS synthesis. #; gene not among the 200 genes shown in Figure 1C and Table S1. Gray boxes with X indicate samples from time courses with insufficient counts at time 0 to assign scores.

#### EPS regulators have a recurrent fitness signature

Insertions in known regulators of EPS production and several other genes had fitness profiles related to, but distinct from, those of most T4P genes. As a group, these mutants were modestly but consistently underrepresented during the first 24 h, moderately overrepresented in non-agg and agg samples at later time points (48-72 h), and strongly underrepresented in starvation-induced spore samples (Fig. 3C; Tables S1 and S2). The Dif chemosensory system regulates EPS production (57) in response to T4P (58). EPS in turn triggers T4P retraction (59), forming a regulatory loop (58) whose molecular basis is incompletely understood (56). Mutations in *difA/C/E*, encoding methyl-accepting chemotaxis protein (MCP), CheW, and CheA homologs, eliminate or severely impair EPS production, T4P-dependent motility, fruiting body development, and cell cohesion (57, 60, 61). Co-development with wild type appears to rescue aggregation of *dif* (aka *dsp*) mutants but not sporulation (62), and adding purified EPS (fibrils) rescues both (57, 63), suggesting EPS can be shared in mixed populations. In our pool, *difA/C/E* (MXAN_6696/6694/6692) insertions were modestly depleted in non-agg and agg samples early in development (Fig. 3C), indicating reduced attachment and/or survival. Later, these mutants were moderately enriched in non-agg and agg samples and strongly depleted in spore samples, indicating that EPS sharing did not rescue their aggregation or sporulation. A *difD* mutant overproduces EPS (64), unlike *difA/C/E* mutants, yet *difD* (MXAN_6693) insertions gave a similar fitness pattern except for no early depletion (Fig. 3C). We predict that insertions in *difD* exert polar effects on *difE* while preserving enough DifE activity to support attachment and/or survival.

EPS production is also influenced by a specialized chaperone system that includes the DnaK homolog SglK (65, 66). The DnaK-GrpE-DnaJ chaperone complex generally responds to heat stress to maintain proteostasis, but in bacteria with complex genomes additional orthologs adopt specialized roles, including regulation of pili and flagella, and coordination of motility machines with polysaccharide biosynthesis (67-70). Strains with insertions in *sglK* (MXAN_6671) and *grpS* (MXAN_6672), homologs of DnaK and GrpE, had fitness profiles resembling *difA/C/E* insertions (Fig. 3C), consistent with defects in EPS production, T4P-dependent motility, aggregation, and sporulation reported previously for *sglK* mutants (65-68). Since *sglK* is co-transcribed downstream of *grpS*, which is itself dispensable for T4P-dependent motility and development (perhaps owing to an unidentified redundant ortholog) (66), the *grpS* pattern likely reflects polar effects on *sglK* (Fig. 3C). DnaK proteins typically act with a DnaJ partner, which was unknown for SglK. Searching for a J-domain protein with a matching fitness pattern, only *sgmC* (MXAN_1106) qualified, with a profile remarkably similar to *sglK* and *grpS* (Fig. 3C). We therefore predict that SgmC partners with SglK and GrpS to regulate EPS production and enable T4P-dependent motility for aggregation and sporulation; this is supported by the known T4P-motility defect of *sgmC* insertions (71).

Insertions in *rasA* (MXAN_4150) gave a profile similar to the regulators above (Fig. 3C), and *rasA* mutants share defects with *difA/C/E* and *sglK* mutants (71, 72). RasA is broadly conserved in Myxococcales and has a DUF2336 domain, but its roles in EPS production, T4P-dependent motility, and fruiting body development remain unclear.

Finally, the HsfA/B two-component system regulates a large number of genes during growth (73) and affects the heat shock response, secondary metabolite production, predation, motility, and development (73-75). Insertions in *hsfA* (MXAN_5364) and *hsfB* (MXAN_5365) gave patterns nearly identical to *difA/C/E*, *sglK*-*grpS*-*sgmC*, and *rasA* (Fig. 3C), suggesting deficient EPS production and/or shared regulation of other processes. Together, these genes define a recurrent fitness signature for known or inferred regulators of EPS production. Because their scores in non-agg and agg samples exceed those of EPS biosynthesis mutants (below), we infer that these regulators modulate more than EPS alone. To explain their near-identical profiles, we propose that HsfA and DifA are substrates of the specialized SglK-GrpS-SgmC chaperone system, given that both interact with SglK (MxDnaK2) in growing cells (67). Experiments with developing cells, and with co-developing mixtures of marked mutant and wild-type strains, can be used to test our proposal.

#### EPS sharing rescues aggregation but not sporulation

The EPS biosynthetic genes lie at the *eps* locus (MXAN_7417-7451), and in-frame deletions in several severely impair aggregation and sporulation in submerged culture (13). In contrast, in our mixed population, insertions in most *eps* genes had <2-fold effect on non-agg and agg fitness scores (Fig. 3C; Tables S1 and S2), indicating little effect on aggregation and implying that EPS sharing rescued aggregation of most *eps* mutants in our mutant pool. Unlike most *eps* mutants, the EPS regulatory mutants showed early attachment and/or survival defects and later aggregation defects that EPS sharing did not rescue (Fig. 3C). Based on these and other differences between *dif* and *eps* mutants previously noted (56), we infer that *dif* and related regulators control functions beyond EPS production. Importantly, *eps* insertions were strongly depleted in starvation-induced spore samples despite this aggregation rescue, as were *dif* mutants (Fig. 3C). This demonstrates that both the Dif system and the EPS pathway are required cell-autonomously for efficient sporulation in the mutant pool.

Several *eps* mutants departed from the majority profile. Insertions in *epsU* (MXAN_7422) and *epsA* (MXAN_7451), both predicted glycosyltransferases, were moderately depleted in non-agg samples, but much less depleted in agg samples (Fig. 3C). This fitness pattern suggests that EPS sharing rescued aggregation of some mutant cells, protecting them from loss, but non-aggregated mutant cells were vulnerable to loss due to poor attachment and/or survival. Insertions in *epsM/L/K/I* (MXAN_7436/7437/7438/7440) were depleted in glycerol-induced spore samples (Fig. 3C). EpsM/L/K likely form an efflux pump important for both sporulation types, and EpsI (Nla24) is a transcriptional activator important for both (76).

Insertions in two operons had fitness patterns similar to the majority of *eps* genes, suggesting a shared function. In-frame *stkA/B* deletion mutants over- or under-produce EPS, respectively, with partial aggregation defects (77). StkA is a DnaK homolog and StkB has a sterol carrier protein 2 (SCP2) domain implicated in lipid transport and metabolism. Insertions in *stkA/B* (MXAN_3474/3475) were enriched 2-fold in 72-h non-agg samples, consistent with mild aggregation defects, and were depleted in starvation-induced spore samples like *eps* mutants (Fig. 3C; Tables S1 and S2). Insertions in the *pglA* (MXAN_7252) and downstream *pkn12* (MXAN_7251) genes, which appear to be co-transcribed, also had fitness profiles resembling most *eps* genes. PglA is a broadly conserved ExoD-family polysaccharide biosynthetic protein whose mutant is partially gliding-defective (78), though whether it acts in LPS O-antigen biosynthesis (important for gliding and development (79)), is unknown. Pkn12 resembles serine/threonine protein kinases (STPKs), but its substrate is unknown. Since a *pkn12* mutant develops normally on starvation agar, both alone and mixed 1:1 with wild type (80), the sporulation defect of *pkn12* insertions in our pool is surprising (Fig. 3C). This indicates that the mixed mutant population does not rescue *pkn12* under submerged culture, and suggests that *pkn12* would also sporulate poorly in monoculture under those conditions.

Insertions in *pglE* (MXAN_5585), MXAN_1943, and the apparent MXAN_1944/1945 operon showed a distinct pattern, with greater enrichment in agg than non-agg samples (Fig. 3C; Table S1), suggesting superior aggregation in the pool despite annotations indicating polysaccharide biosynthesis. PglE is a predicted transmembrane glycosyltransferase whose mutant has 3-fold reduced gliding probability (78). Like PglA, it is not known whether PglE is involved in LPS biosynthesis. MXAN_1943 is a predicted cytoplasmic oxidoreductase. MXAN_1944/1945 are predicted cytoplasmic glycosyltransferase/polysaccharide deacetylase enzymes. None of these genes lie near known cell-surface-polysaccharide clusters (56). Determining whether their products modify known or novel polysaccharides will be important, since our results suggest that loss of particular modifications can enhance the aggregation of individual cells in a mixed population.

#### SCP biosynthesis genes and new sporulation factors

SCP forms a protective layer around both starvation- and glycerol-induced spores (1, 56) (Fig. 3A). Three genomic clusters encode Exo proteins that synthesize, modify, and export the coat polysaccharide chains (26, 81, 82). A fourth cluster encodes Nfs proteins that appear to form a cell-envelope complex (5, 26) that shortens these chains for assembly into a stress-resistant coat (81). As expected, insertions in most genes of the *exoA-I* (MXAN_3225-3233), *exoJ/K* (MXAN_3026/3027), *exoL-P* (MXAN_3259-3263), and *nfsA-H* (MXAN_3371-3378) clusters were strongly depleted in both spore sample types (Fig. 1 and 3C; Tables S1 and S2) while remaining largely unaffected in non-agg and agg samples, consistent with the normal aggregation reported for defined mutants (5, 26, 82). Insertions in *nla6* (MXAN_4042), encoding a direct transcriptional activator of the *exoA-I*, *exoL-P*, and *nfsA-H* operons (83, 84), had a similar fitness pattern (Fig. 3C; Table S1) as expected from studies of a defined *nla6* mutant (76). Insertions in *wecB* (MXAN_1101) were strongly depleted in both types of spore samples (Fig. 3C; Table S2). WecB resembles bacterial epimerases involved in cell-surface polysaccharide biosynthesis and is required in *M. xanthus* for coat material to accumulate at the spore surface (26). Insertions in genes for a predicted glycosyltransferase (MXAN_3469) and a GlmS sugar transaminase (MXAN_1386) were likewise depleted in both spore types (Fig. 3C; Table S1). All of these proteins likely participate in SCP synthesis, as do the glycosyltransferases and other sugar-modifying enzymes encoded in the *exo* and *nfs* clusters and by *wecB* (26, 81, 82).

Insertions in several genes not previously linked to SCP synthesis were also notably depleted in both spore-sample types (Fig. 3C; Tables S1 and S2). Insertions in the response regulator (RR) gene MXAN_3738 were enriched in non-agg samples, consistent with a moderate aggregation defect, as were insertions in the apparent MXAN_4674-4672 operon. MXAN_3738 orthologs are broadly conserved in Myxococcales and have a typical RR N-terminal receiver domain but an atypical C-terminal domain with an unusual predicted β-sheet fold. MXAN_4674-4672 encode MaoC-family dehydratases and a reductase, which may act in fatty acid oxidation and cell wall remodeling, respectively, processes important for sporulation (33, 53). Figure 3C also includes three *mnm* genes (MXAN_7494, 7506, 3499) and *cydA* (MXAN_6913). The *mnm* enzymes modify tRNAs at position 34, presumably affecting codon-anticodon recognition, and *cydA* encodes a terminal oxidase of the aerobic respiratory chain. Insertions in *cydA* and *mnmA* were depleted in both non-agg and agg samples, indicating poor attachment and/or survival. Together, these results confirm the importance of *exo*, *nfs*, *nla6*, and *wecB* for both sporulation pathways and uncover new genes affecting both, including an atypical RR and enzymes potentially involved in SCP formation, cell-envelope remodeling, and the metabolic changes underlying the rod-to-spore transition.

#### LPS biosynthesis genes affect cell loss and sporulation

LPS is composed of lipid A embedded in the OM surface leaflet, a core oligosaccharide attached to lipid A, and a variable O-antigen polysaccharide attached to the core. *M. xanthus* LPS O-antigen is important for gliding motility and fruiting body development (79, 85). In a model for biosynthesis of the O-antigen, WbgA (MXAN_4621) and WbgB (MXAN_4619) are glycosyltransferases, and SgmR (MXAN_4620) is a methyltransferase, forming O-antigen chains transported by Wzm (MXAN_4623) and Wzt (MXAN_4622) across the inner membrane (79). We observed that insertions in *wbgB*, *sgmR*, and *wzt* were moderately underrepresented in non-agg samples and less so in agg samples, similar to insertions in *epsU* and *epsA* (Fig. 3C; Tables S1 and S2). We propose that LPS sharing *via* OME rescued aggregation of some mutant cells, protecting them from loss, but non-aggregated mutant cells were vulnerable to loss due to poor attachment and/or survival. Developmental sharing of LPS *via* OME partially rescued sporulation of a *wzm* mutant in a 1:1 mixture with wild type (86). LPS sharing, like EPS sharing, did not completely rescue starvation-induced sporulation in our mutant pool (Fig. 3C). Insertions in *wbgB* and *sgmR* were ∼2-fold enriched in glycerol-induced spore samples, a result warranting further investigation. Insertions in MXAN_4614 and *pglF* (MXAN_4616) resembled those in *wbgB*, *sgmR*, and *wzt*, which together with the proximity of these genes suggests roles in LPS production. A *pglF* mutant is partially gliding-defective, and PglF is a predicted cytoplasmic glycosyltransferase implicated in polysaccharide synthesis (78). *pglF* does not appear to be co-transcribed with the upstream MXAN_4614, also a predicted cytoplasmic glycosyltransferase. To our knowledge, an MXAN_4614 mutant has not been created previously.

### Cell loss early in development can be influenced by many genes

Mutants depleted in non-agg and agg samples early in starvation-induced development identify genes important for attachment and/or survival (Fig. 1 and 4). Diminished function of these genes in wild type may contribute to cell death. About 90% of cells die under our submerged-culture conditions (9), but the mechanism(s) of death remain unclear. A mechanism involving a toxin-antitoxin system contributes to death during development of a mutant with a compromised OM secretin (87), but does not account for developmental lysis of wild type (7, 88). Having already described several genes important for attachment and/or survival (Fig. 2B and 3C), we now highlight additional classes of such genes.

**FIG 4.**
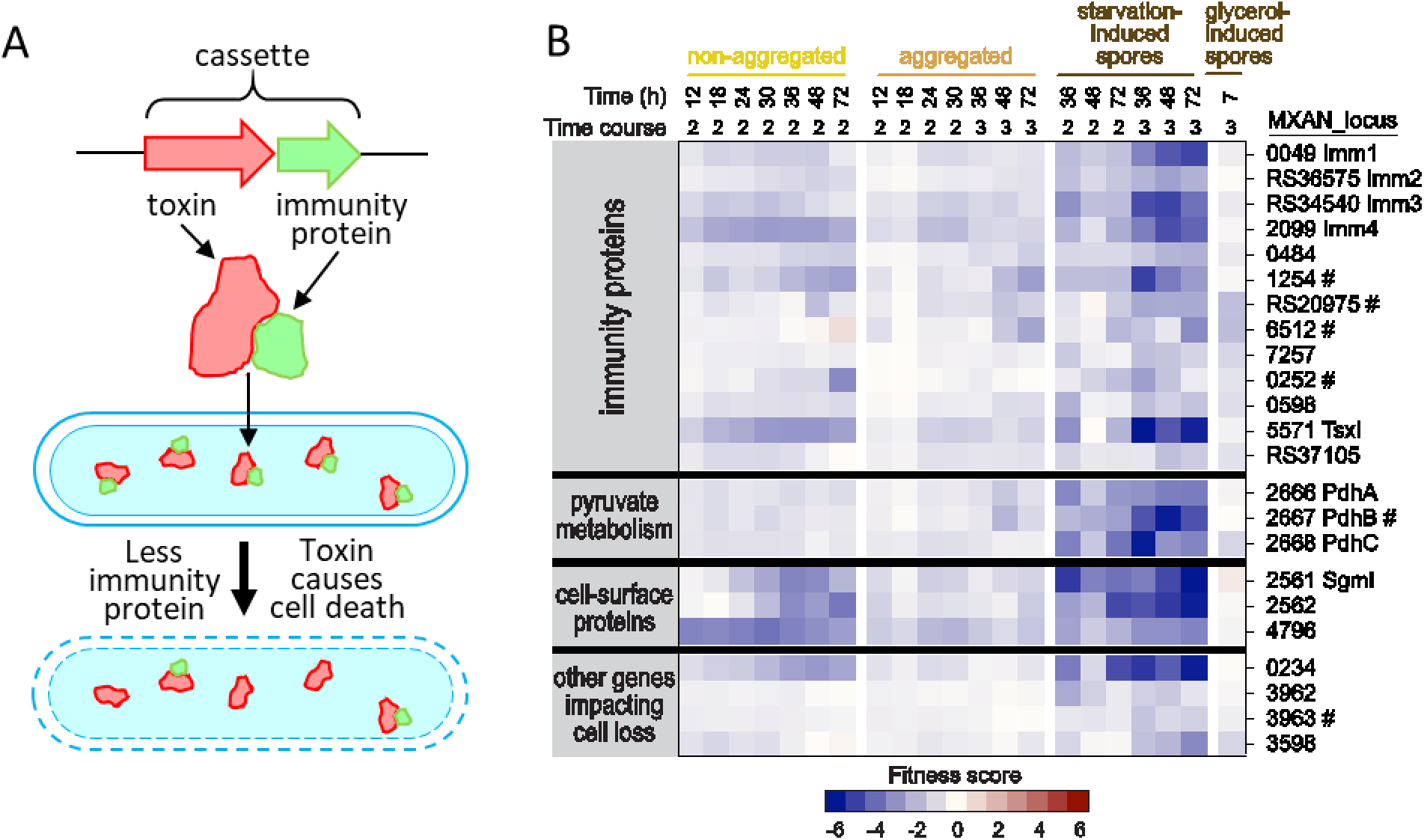
Cell survival during development depends on toxin-immunity cassettes and other genes. (A) Typical toxin-immunity cassette. Upstream toxin gene co-transcribed with gene for immunity protein. Immunity protein binds to toxin in cells that survive. Toxin kills developing cell if immunity protein level is too low. (B) Heat map showing fitness scores of genes important for developmental survival. #; gene not among the 200 genes shown in Figure 1C and Table S1.

#### Loss of toxin immunity is a frequent cause of cell loss

*M. xanthus* has numerous mechanisms for predation and kin discrimination (1, 89), and this abundant killing potential carries a risk of self-harm. Immunity proteins protect cells from the toxin proteins used in kin discrimination. A toxin is typically encoded upstream of and adjacent to its cognate immunity protein, forming a toxin-immunity cassette (Fig. 4A); too little immunity protein can lead to toxin-mediated death. Crosstalk between sufficiently similar cassettes allows discrimination between close and distant relatives (90), and explains the unexpected viability of some immunity-gene insertion mutants (80). Insertions in *imm1/2/3/4* (MXAN_0049/RS36575/RS34540/2099) produced mutants depleted in starved samples (Fig. 4B; Table S1), consistent with the poor sporulation of *imm1/3/4* mutants co-developed with wild type (*imm2* has not been previously tested) (80). Imm1/2/3/4 are Imm11-family proteins, and their cognate Tox1/2/3/4 are AHH/ENDO VII nucleases delivered to neighboring cells by the type VI secretion system (T6SS) (90, 91). Among the 10 other *imm11*-family genes, insertions in MXAN_0484/1254/RS20975 have fitness scores indicative of poor attachment and/or survival (Fig. 4B; Tables S1 and S2). These genes encode SitI5-family immunity proteins whose cognate AHH toxins are SitA5-family lipoproteins transferred by OME (92).

The *M. xanthus* wild-type strain we used (DK1622) also encodes non-AHH kin-discrimination toxins, including 15 SitA toxins transferred by OME (92) and two toxins (TsaE and TsxE) delivered by the T6SS (93, 94). Among the cognate immunity genes, insertions in four *sitI*-family genes (MXAN_6512/7257/0252/0598) and in *tsxI* (MXAN_5571) resulted in poor attachment and/or survival (Fig. 4B; Tables S1 and S2).

Studies of natural *M. xanthus* isolates implicated a 150 kb polymorphic region in kin discrimination (95) and showed that rearrangement hotspot (Rhs) proteins mediate the effect (94). Strain DK1622 encodes two Rhs proteins. One (MXAN_5799) lies upstream of and adjacent to MXAN_RS37105; insertions in MXAN_RS37105 resulted in poor attachment and/or survival (Fig. 4B; Table S1), suggesting that MXAN_5799-RS37105 is a toxin-immunity cassette.

Insertions in some immunity genes mentioned above had distinct fitness effects. Insertions in *imm4* and *tsxI* were more depleted in non-agg than agg samples, suggesting different toxin/immunity protein ratios in these populations due to cell autonomous (e.g., proteolysis) and/or non-autonomous (e.g., toxin transfer) differences. Insertions in several *sitI* genes and MXAN_RS37105 were 2- to 4-fold depleted in glycerol-induced spore samples, indicating poor survival and/or sporulation.

#### Metabolic, cell-surface, and cytoplasmic genes also support attachment and/or survival

The pyruvate dehydrogenase (Pdh) complex is critical for energy generation during growth, but its role in development is unclear. *pdhA*/*B*/*C* (MXAN_2666/2667/2668) mutants were depleted in starved samples (Fig. 4B; Tables S1 and S2), indicating that PdhA/B/C are important for attachment and/or survival as early as 12 h into development and are particularly important for sporulation.

Bacterial fibronectin type III domain-containing proteins are cell-surface proteins that typically mediate adhesion to host cells, other bacterial cells, the extracellular matrix, or a polysaccharide substrate. One such *M. xanthus* protein, SgmI (MXAN_2561), is implicated in T4P-dependent motility (71), but is poorly characterized. SgmI has two central predicted fibronectin type III (immunoglobulin-like) domains and a C-terminal sorting tag presumed to direct it to the cell surface for anchoring *via* the type II secretion system (29, 96). MXAN_2562 lies downstream of and appears to be co-transcribed with *sgmI* and encodes a predicted OM β-barrel protein, while MXAN_4796 is an uncharacterized predicted OM lipoprotein with four surface-exposed fibronectin type III domains. Insertions in *sgmI*, MXAN_2562, and MXAN_4796 had fitness profiles (Fig. 4B; Table S1) resembling insertions in *epsU*, *epsA*, and known or implied LPS biosynthesis genes (Fig. 3C). We propose that SgmI, MXAN_2562, and MXAN_4796 sharing *via* OME, like sharing of LPS *via* OME and EPS *via* secretion, rescued aggregation of some mutant cells, protecting them from loss, but non-aggregated mutant cells were vulnerable to loss, probably due to poor attachment of cells lacking these cell-surface adhesion proteins, and starvation-induced sporulation was not rescued completely.

A few proteins with predicted cytoplasmic functions also affected developmental attachment and/or survival. These include the putative anti-σ factor MXAN_0234, the iron-sulfur cluster-binding protein MXAN_3962, the glycine/D-amino acid oxidase MXAN_3963, and the DNA cytosine methyltransferase MXAN_3598 (Fig. 4B; Tables S1 and S2; see SI for more discussion). Among these, insertions in MXAN_0234 were more depleted in non-agg than agg samples, likely stemming from differential gene expression in the populations.

Collectively, these results show that many genes can contribute to cell loss during *M. xanthus* development. We propose that diminished function of one or more (possibly many) of these genes in starving cells accounts for the massive population death normally observed during fruiting body development.

### Developmental winners and a connection between cheating and phase variation

Our analysis identified a large group of insertion mutants that were overrepresented in starvation-induced spore samples, which we call developmental winners (Fig. 1 and 5A). Three types of developmental winners have been described in previous studies of co-developed *M. xanthus* strains: *1)* cheaters benefit from a cooperative social interaction without contributing to its cost (97), and sporulate poorly in monoculture; *2)* facultative social exploiters benefit by exploiting a strain of different genotype while retaining all normal social interactions (98), so they sporulate normally in monoculture; *3)* a superior cooperator sporulated better than wild type in monoculture (99), by losing a small RNA (Pxr-S) that negatively regulates entry into development (100), and affects both social interactions and gene expression (99, 101).

**FIG 5.**
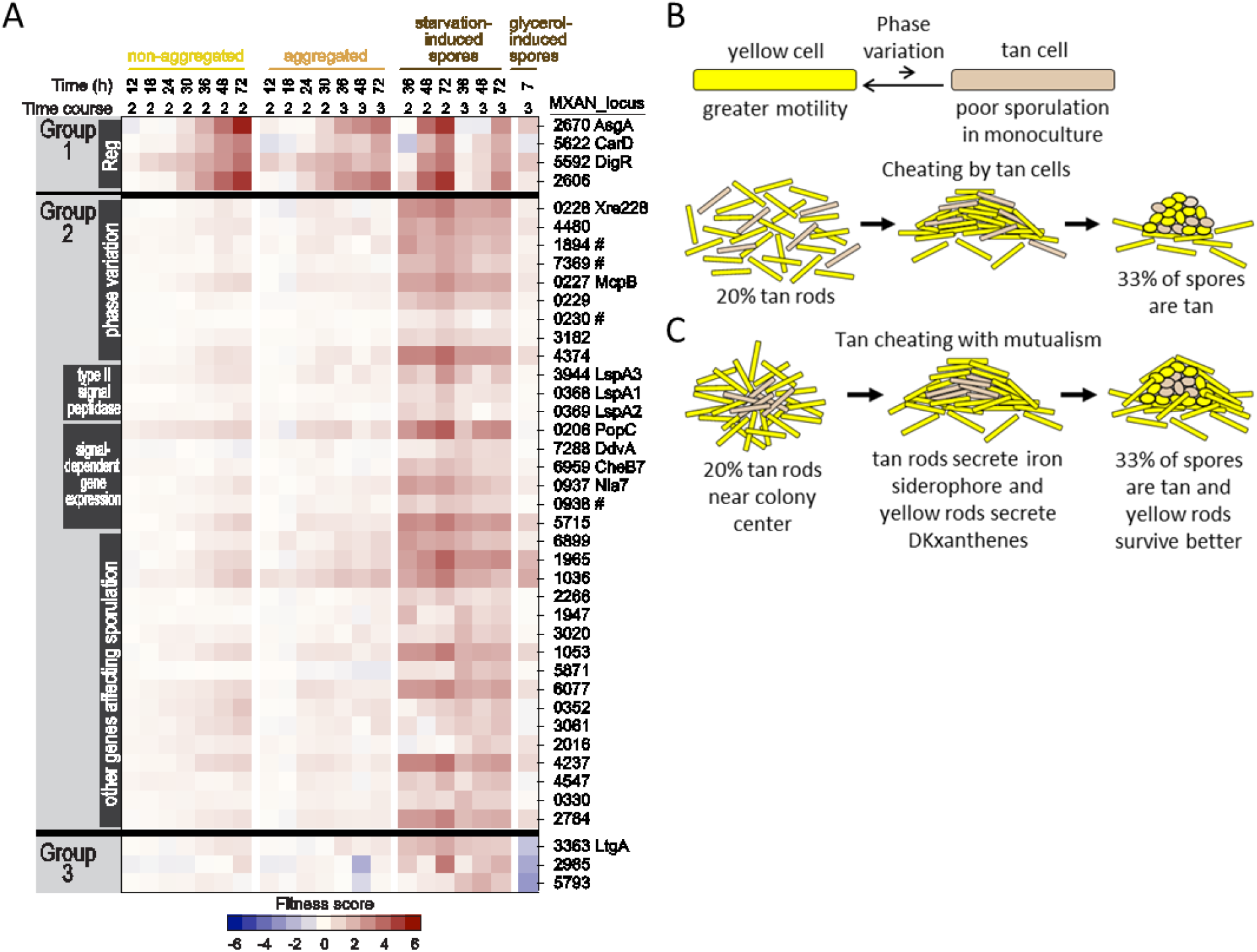
Developmental winners, cheating, and mutualism. (A) Heat map showing fitness scores of insertion mutants with increased starvation-induced sporulation. #; gene not among the 200 genes shown in Figure 1C and Table S1. (B) Phase variation favors yellow variants with greater motility and better sporulation in monoculture than tan variants, but in a mixture of rods randomly placed on a surface, tan cells cheat on a signal produced by yellow cells and form spores more efficiently. (C) Mutualism model. See text.

Insertions in the genes listed in Figure 5A may reveal new types of developmental winners. For example, insertion in a gene encoding a negative regulator that acts only cell-autonomously (not socially) would represent a different type of winner than the superior cooperator, although both would sporulate better than wild type in monoculture. For several genes in Figure 5A, a defined mutant was previously shown to sporulate poorly in monoculture relative to wild type (see below). Taken together with our results, we conclude that these mutants are cheaters; some are tan phase variants.

*M. xanthus* exhibits phase variation in colony color and morphology that correlates with the ability to form spores in mixed populations. The predominant colony type is yellow with a rough surface and a jagged edge indicative of greater motility (Fig. 5B), whereas a minority are tan with a smooth surface and edge. A yellow-colony inoculum grows to a population of ∼95-99% yellow and ∼1-5% tan cells, while a tan-colony inoculum grows to ∼75% tan and ∼25% yellow cells (102). In co-developed mixtures, tan cells preferentially form spores (Fig. 5B), suggesting a connection between the phase of a cell upon entering development and its ability to sporulate (103). Since tan phase variants typically form fewer spores in monoculture (103-105) (Fig. 5B), tan variants may be cheaters, and mutations that influence phase variation may cause cheating. Below we describe two groups of developmental winners connected to phase variation and cheating, followed by a third group with a distinct fitness profile.

#### A first group of regulatory mutants cheats during starvation-induced development

Strains with insertions in the regulatory genes *asgA* (MXAN_2670), *carD* (MXAN_5622), *digR* (MXAN_5592), and MXAN_2606 became increasingly enriched in non-agg, agg, and starvation-induced spore samples over time (Fig. 5A; Table S1), indicating poor aggregation but enhanced sporulation. Defined *asgA*, *carD*, and *digR* mutants in monoculture have severe aggregation defects (106-108), consistent with our results, but they also have severe sporulation defects (106-108). The spore enrichment we observed therefore indicates that *asgA*, *carD*, and *digR* insertions enabled cheating in our Tn-Himar mutant pool.

AsgA functions in a regulatory pathway with AsgB to produce extracellular A-signal (106). We did not obtain fitness scores for *asgB* (MXAN_2913), but a defined *asgB* mutant was previously shown to be a cheater (97). AsgB is likely a transcription factor (109) and AsgA is a hybrid RR-histidine kinase (HK) (110). Both proteins regulate expression of secreted proteases that may produce A-signal (111), a mixture of amino acids and peptides that serves as a quorum-sensing signal early in development (112). The *asgA/B* mutants presumably cheat by avoiding the cost of A-signal production while benefiting from A-signal made by the mixed population. CarD is a global regulator of transcription (113). Defined *asgA* and *carD* mutants were partially rescued by co-development with wild type at an initial ratio of 1:1, but cheating was not observed (106, 108). Similarly, the defined *asgB* mutant did not cheat when mixed 1:1 with wild type, but cheated when mixed with a 100-fold excess of wild type (97). The small proportion of *asgA*, *carD*, and *digR* mutants in our pool may similarly permit cheating, a hypothesis that can be tested by co-developing defined mutants with excess wild type.

DigR is an RR that regulates extracellular matrix (ECM) composition (107), so *digR* mutants may cheat on ECM produced by a mixed population. DigR contains a helix-turn-helix (HTH) DNA-binding domain of the xenobiotic response element (Xre) superfamily (107). Other transcription factors in this superfamily (see below) have been proposed to form a regulatory hierarchy governing *M. xanthus* phase variation (104); consistent with a phase-variation link, defined *digR* and *asgA/B* mutants are tan phase variants (106, 107). MXAN_2606 is an uncharacterized orphan hybrid HK-RR-HK. If a defined mutant proves to have a monoculture sporulation defect during starvation-induced development, the spore enrichment we observed (Fig. 5A) would indicate that MXAN_2606 insertion mutants also cheated in our pool. Insertions in *asgA*, *carD*, *digR*, and MXAN_2606 still formed glycerol-induced spores (Fig. 5A), consistent with a result for a defined *carD* mutant (108). Other defined mutants remain to be tested.

#### A second mutant group links phase variation to enhanced sporulation

A group of developmental winners exhibited weakly increasing positive fitness scores over time in non-agg and agg samples (or scores near zero), combined with positive fitness scores in starvation-induced spore samples. Several members are disrupted in HTH-Xre regulators: Xre228 (MXAN_0228) governs phase variation (114) and MXAN_4480, MXAN_1894, and MXAN_7369 are proposed regulators of phase variation (104) (Fig. 5A; Tables S1 and S2). Insertions in genes near *xre228* also gave positive scores in spore samples, including *mcpB* (MXAN_0227), which encodes an orphan MCP (115), and MXAN_0229 and MXAN_0230, which encode hybrid RR-HK-RR and HK-RR-RR proteins, respectively. This suite of genes is conserved and syntenic across multiple myxobacteria (Fig. S2). An *xre228* mutant is a tan phase variant (114), and tan variants are deficient in the yellow DKxanthene pigments produced from the *dkx* biosynthetic cluster (105). Notably, three tan variants (*xre228*, *dkxG*, and *asgB*) overexpress iron-acquisition genes relative to a yellow wild type (104), prompting a proposed mutualism: tan cells, being less motile, accumulate in colony centers and within developing mounds where their enhanced iron siderophore production benefits surrounding yellow cells’ survival (Fig. 5C). In turn, DKxanthenes produced by yellow cells would stimulate preferential sporulation of tan cells. This yellow-tan mutualism was proposed to be critical for *M. xanthus* survival in nature (104). The same authors proposed that gene clusters encoding STPKs and DUF2381-family proteins regulate phase variation. Consistent with this, insertions in two genes within such clusters (MXAN_3182, MXAN_4374) were overrepresented in spore samples (Fig. 5A; Table S1), supporting a tan-phase sporulation bias in mixed populations with yellow cells (103), and possibly meeting the criteria for cheating if defined mutants prove sporulation-defective.

Not all cheaters are tan variants, judging by the colony color of defined mutants. The defined *carD* mutant noted above is yellow (108). Insertions in three type II signal peptidase genes, *lspA3* (MXAN_3944), *lspA1* (MXAN_0368), and *lspA2* (MXAN_0369), were overrepresented in starvation-induced spore samples (Fig. 5A; Table S1). Only the defined *lspA3* mutant is a tan variant with delayed fruiting body formation, whereas *lspA1* and *lspA2* mutants are yellow with normal fruiting body formation (116). The second group of developmental winners also includes genes whose defined mutants were characterized and presumably yellow, since none were reported as tan variants: *mcpB* (115), *popC* (MXAN_0206) (117), *ddvA* (MXAN_7288) (113), *cheB7* (MXAN_6959) (118), and *nla7* (MXAN_0937) (76) (Fig. 5A; Table S1) are all involved in signal-dependent gene expression (SI). Finally, the second group includes 18 other genes without reported defined mutants, encoding proteins of diverse predicted function (Fig. 5A; Tables S1 and S2; see SI for more discussion of these genes).

#### A third group has opposite effects on the two sporulation pathways

A group of developmental winners with insertions in *ltgA* (MXAN_3363), MXAN_2985, and MXAN_5793 had negligible fitness scores in non-agg and agg samples, positive scores in starvation-induced spore samples, and negative scores in glycerol-induced spore samples (Fig. 5A; Table S1). Consistent with our results, a defined *ltgA* mutant aggregates normally upon starvation, but is defective in glycerol-induced sporulation (33). LtgA is a lytic transglycosylase that degrades PG. MXAN_2985 is uncharacterized. MXAN_5793 affects Pxr-S function (119) (SI; Fig. S3). Understanding how these proteins exert opposite effects on starvation-versus glycerol-induced sporulation efficiency is an intriguing topic for future studies (see SI for additional discussion).

Our results show that numerous insertion mutants become enriched in starvation-induced spore samples during a single round of development by the Tn-Himar pool. Creating defined mutants and examining their monoculture development can distinguish among the possible explanations for these winners. Of the 41 genes we identified (Fig. 5A), defined mutants have been reported for 13, and seven of these had impaired monoculture development (*asgA*, *digR*, *carD*, *xre228*, *lspA3*, *popC*, *ddvA*) (106-108, 113, 114, 116, 117), indicating that insertions in those seven enabled cheating in our mixed population (Fig. 5A). None of the seven defined mutants were known cheaters, but four (*asgA*, *digR*, *xre228*, *lspA3*) (106, 107, 114, 116) were known tan variants, supporting a proposed connection between tan variants and cheating in mixed populations (103, 104) (Fig. 5B and 5C). The six other defined mutants aggregate normally (*lspA1*, *lspA2*, *nla7*, *ltgA*) (33, 76, 116), prematurely (*mcpB*) (115), or with delay (*cheB7*) in monoculture, but their sporulation was not quantified except for the *nla7* and *cheB7* mutants, which were normal (76, 118). This implies that *nla7* and *cheB7* insertion mutants exhibited facultative social exploitation in our Tn-Himar mutant pool (98). Quantitative sporulation assays of the other four defined mutants, and of defined mutants for the remaining 28 genes (Fig. 5A), will reveal the relative contributions of facultative social exploitation, cheating, and cell-autonomous mechanisms to spore enrichment. Developmental winners have been observed in experimental evolution studies (97, 99, 120), but evolved strains typically carry multiple mutations, and only once has the gene responsible for enhanced sporulation been identified (99). Our approach powerfully expedites the discovery and analysis of developmental winners.

### Cell envelope, CRISPR-Cas, and regulatory genes primarily important for sporulation

A group of insertion mutants was strongly depleted in one or the other type of spore sample, with weaker or no effects in non-agg and agg samples (Fig. 1 and 6). Most of the corresponding genes are uncharacterized or poorly characterized with respect to development, and bioinformatic analysis suggests roles in a variety of cell-envelope functions. The exceptions are the *devT/R/S* genes, part of a CRISPR-Cas locus known to be important for starvation-induced sporulation, and *askB/C*, which encode signal transduction proteins critical for glycerol-induced sporulation. Most genes in this group offer new opportunities to explore the cell-envelope functions underlying spore formation.

**FIG 6.**
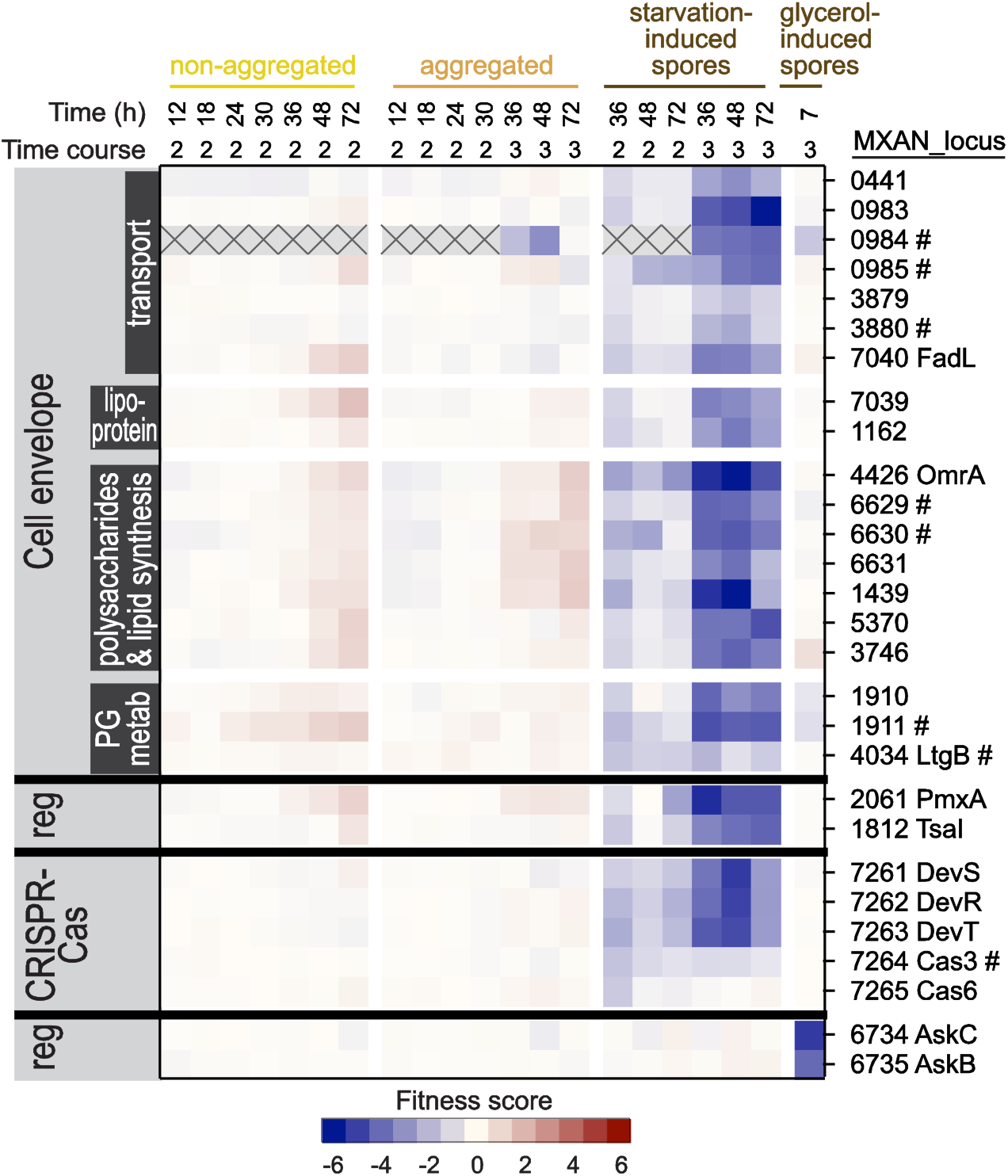
Cell envelope, CRISPR-Cas, and regulatory genes important for one or the other type of sporulation. Heat map showing fitness scores of insertion mutants with decreased starvation- or glycerol-induced sporulation. #; gene not among the 200 genes shown in Figure 1C and Table S1. Gray boxes with X indicate samples from time courses with insufficient counts at time 0 to assign scores.

#### Uncharacterized cell-envelope genes are required mainly for fruiting body sporulation

Insertions in a group of developmentally uncharacterized cell-envelope genes were depleted in starvation-induced spore samples, with little or no effect on non-agg and agg samples (Fig. 6; Tables S1 and S2). These genes encode predicted transporters (MXAN_0441, 0983/0984/0985, 3880), lipoproteins (MXAN_1162, 7039), *omrA* (MXAN_4426) encoding a putative phospholipid flippase (121), and predicted enzymes for lipoquinone biosynthesis (MXAN_6629) and adjacent oxidoreductases (MXAN_6630/6631), glycosylation (MXAN_1439, 5370), oxidoreduction (MXAN_3746), PG recycling (MXAN_1910), and β-lactam inactivation (MXAN_1911). A predicted hybrid HK-RR-RR (MXAN_3879) is encoded adjacent to one of the transporters above and exhibits a similar fitness profile. MXAN_7039 appears to be co-transcribed with downstream *fadL* (MXAN_7040) encoding a fatty acid uptake transporter (53), and some of the other cell-envelope genes lie in apparent operons (Fig. 6; see SI for additional discussion of these genes).

LtgB (MXAN_4034) is a lytic transglycosylase important for PG degradation during starvation-induced spore formation that also inhibits LtgA binding to PG during glycerol-induced sporulation (33). Consistent with this starvation-induced role, insertions in *ltgB* were depleted in starvation-induced but not glycerol-induced spore samples (Fig. 6).

Insertions in regulatory genes *pmxA* (MXAN_2061) and *tsaI* (MXAN_1812) resulted in similar fitness profiles as insertions in *ltgB* and the uncharacterized cell-envelope genes (Fig. 6). PmxA is a phosphodiesterase that preferentially degrades cyclic GMP-AMP (cGAMP) (122). A *pmxA* mutant fails to aggregate and shows 6-fold reduced sporulation (123). In contrast, *pmxA* insertions yielded only ∼2-fold enrichment in 72-h non-agg samples but 20- to 30-fold depletion in time course 3 starvation-induced spore samples (Fig. 6; Table S1). We conclude that aggregation of *pmxA* mutants is largely rescued in our Tn-Himar pool, while *pmxA* is required cell-autonomously for efficient sporulation, presumably to lower the cGAMP level (124). TsaI is the immunity protein of a toxin-immunity cassette that depends on the T6SS for TsaE (MXAN_1813) toxin delivery (94, 125). Insertions in *tsaI* had little effect on non-agg and agg samples (Fig. 6), indicating normal attachment and/or survival, unlike the effects of insertions in many other toxin immunity genes (Fig. 4). TsaE is also unusual; it has antifungal activity (125). TsaE also appears to have mild toxicity to *M. xanthus* since a *tsaI* mutant was slightly outcompeted by wild type during co-incubation on nutrient agar. The strong depletion of *tsaI* insertions in fruiting body spore samples (Fig. 6) implies that in the absence of immunity, TsaE is more toxic to *M. xanthus* during development than during growth. Collectively, these results reveal novel cell-envelope functions in transport, lipid and polysaccharide synthesis, PG metabolism, and regulatory proteins that act primarily in sporulation during fruiting body development.

#### The dev CRISPR-Cas locus is required for starvation-induced sporulation

The *dev* operon was among the first CRISPR-Cas loci linked to gene regulation and bacterial development (126). *devT/R/S* (MXAN_7263/7262/7261) insertional mutants were strongly underrepresented in starvation-induced spore samples with little or no effect on non-agg and agg samples (Fig. 6; Table S1), consistent with the normal aggregation and poor sporulation of in-frame deletion mutants (127, 128). The *dev* operon includes upstream *cas6* (MXAN_7265) and *cas3* (MXAN_7264) genes (126), whose in-frame deletions reduce sporulation, though less than *devT/R/S* deletions do (127, 128), again matching our results (Fig. 6; Tables S1 and S2).

#### AskB and AskC are regulatory proteins required for glycerol-induced sporulation

AskB and AskC are hybrid HK-RR-RR-RR and RR-HK proteins, respectively, involved in the responses of *M. xanthus* to glycerol and to ambruticin VS-3, a compound produced by the myxobacterium *Sorangium cellulosum* (4). Strains with insertions in *askC/B* (MXAN_6734/6735) were depleted in glycerol-induced spore samples and largely unaffected in all other samples (Fig. 6; Table S1). Our results are consistent with previous work showing that in-frame deletion of *askC* impairs glycerol-induced sporulation, but not starvation-induced aggregation and sporulation (4). However, in that work, deletion of *askB* alone or together with *askC* did not impair glycerol-induced sporulation, so our finding that *askB* insertions impair this process is unexpected and merits further investigation.

### Concluding remarks

By tracking a barcoded transposon mutant population through time as a mixed pool, we resolved how the loss of individual genes affects the developmental fate of cells within a genetically diverse community, distinguishing requirements for attachment and/or survival, aggregation, and distinct routes of sporulation. This fate-resolved, mixed-population view recovered the expected behavior of many characterized mutants and uncovered numerous new ones, and it separated cell-autonomous functions from those that neighboring cells can supply. Above all, the pooled screen exposed the social dimension of development directly: it identified genes whose loss compromises attachment and/or survival amid the extensive cell death that accompanies development, and it revealed developmental winners that sporulate more efficiently than their neighbors, connecting gene function to cheating and phase variation in some cases.

Our results also refine understanding of how *M. xanthus* motility contributes to development. Although *M. xanthus* motility systems have long been known to participate in development, the relative contributions of T4P-dependent and gliding motility to aggregation versus sporulation have not been thoroughly explored. Our measurements showed that T4P-dependent motility has a substantially larger effect than gliding motility on both processes, and identified new genes likely affecting each system. Our results also bear on the role of OME in *M. xanthus* biology. The OM lipoproteins Tgl and CglB/C/E are transferred between cells by OME during growth, yet in our experiments insertions in their genes behaved like the broader sets of T4P and gliding mutants, indicating that OME-mediated sharing did not rescue development in our pool. On the other hand, it is established that OME can greatly influence the development of mixed cultures (29); OME is one of three kin-discrimination systems (94) that presumably help determine developmental outcomes in natural populations (89).

The approach we describe is readily extended beyond the processes examined here. Our protocol selected for sonication-resistant spores arising from the mixed mutant library, but straightforward modifications could probe other biological processes, including stress responses, survival, and germination. Glycerol-induced spores could be challenged with heat, desiccation, UV irradiation, or antimicrobials, and starvation-induced spores could be dispersed from fruiting bodies by sonication before similar challenges. Proteins important for germination and outgrowth could be identified by shaking spores in nutrient medium long enough for sufficient DNA to be extracted from the resulting vegetative rods.

More broadly, the *M. xanthus* library we have created constitutes a reusable platform for connecting genes to fitness in mixed populations. Such populations can be subjected to repeated cycles of feast and famine, allowing the genetic basis of developmental success to be followed over experimental-evolution timescales. The gene functions catalogued here thus become a starting point for asking how individual genotypes rise, persist, or are lost within the social communities in which *M. xanthus* develops in natural ecosystems.

## METHODS

### Construction of a barcoded *M. xanthus* Himar transposon mutant library

Barcoded Himar transposons were introduced into *M. xanthus* DK1622 (129) *via* conjugation with *Escherichia coli* donor strain WM3064 harboring the pKMW3 mariner transposon vector library (APA752) (6) (see SI for details). Briefly, a DK1622 starter culture was grown overnight, then the culture was diluted into fresh nutrient medium and grown to late-log phase. Simultaneously, an *E. coli* APA752 starter culture was grown overnight in nutrient medium supplemented with diaminopimelic acid (DAP), then the culture was diluted into fresh medium with DAP and grown until mid-log phase. *E. coli* and *M. xanthus* cells were harvested, concentrated cell suspensions were mixed to achieve a 10:1 ratio of donor:recipient cells, and the mixture was spotted on nutrient agar with DAP to allow conjugation. The cell mixture was scraped into liquid medium, cells were suspended, a small portion was used to determine the mutation efficiency, and the remaining conjugation mixture was adjusted to 10% glycerol and stored in aliquots at −80°C. Based on the mutation efficiency, several conjugation experiments were performed to generate a pooled library of approximately 200,000 Tn-Himar insertion mutants.

### Mapping Tn-Himar insertion sites in the *M. xanthus* mutant library

Tn insertion sites were mapped and correlated to barcodes using a nested two-step PCR approach based on the approach outlined by Wetmore *et al.* (6) and described in (130) with minor additional modifications (SI). Tn-Himar insertion mapping data are available at the NCBI Sequence Read Archive under BioProject accession PRJNA1517256.

### Growth and development of the barcoded Tn-Himar mutant library

The *M. xanthus* mutant library was grown in nutrient medium and harvested cells were suspended in starvation buffer and placed in plastic wells to initiate fruiting body development in submerged culture as described (8) (see SI for details). Briefly, a starter culture of the mutant library was grown overnight, then the culture was diluted into fresh medium and grown to the mid-log phase. Cells were harvested, suspended in buffer, and added to buffer in 6-well plastic plates. The plates were incubated and samples were collected at indicated times. To induce sporulation chemically, glycerol was added to culture in the mid-log phase (3) (SI). Two biological replicates of each experiment were conducted, each with three technical replicates.

### Separation of cells with different developmental fates

Low-speed centrifugation was used to separate non-agg from agg cells (7) and we devised a method to isolate spore DNA (see SI for details). Briefly, to separate non-agg from agg cells, cell pellets of developmental samples were suspended in buffer, centrifuged at low speed, the supernatant containing non-agg cells was transferred to a different tube, both tubes were centrifuged at higher speed to pellet cells, the supernatants were removed, and the pellets of non-agg and agg cells were stored at −80°C. To isolate spore DNA from additional developmental samples, agg cell pellets from starvation-induced sporulation and cell pellets from glycerol-induced sporulation were suspended in buffer, sonicated to lyse non-spore cells, treated with DNase I to digest non-spore DNA and then with EDTA to inactivate DNase I, and finally beaten with glass beads to release spore DNA.

### Amplification, sequencing and analysis of Tn-Himar barcoded strains

To assess barcode abundances, we amplified barcodes using the PCR approach developed by Wetmore *et al.* (6). Barcode amplicons were sequenced (SeqCenter, Pittsburg) and analyzed using custom scripts developed by Wetmore *et al.* (6) to calculate gene fitness scores and t-scores, see (SI) and (6) for more details. Barcode amplicon sequencing data for all experiments are available at the NCBI Sequence Read Archive under BioProject accession PRJNA1517256. The individual fitness- and t-scores for each sample are compiled in Table S3 (see Table S3 at [https://doi.org/10.6084/m9.figshare.33346104]). The scores of the technical replicates in each biological replicate were averaged. Genes with an absolute value of an average t-score >4 in at least one biological replicate sample were identified. Among those genes, those with an absolute average fitness score >1 in at least one complete sample (i.e., 2 biological replicates, each with 3 technical replicates) were analyzed further (Table S1; see SI for details).

### Predicted operons and protein transmembrane regions and functions

Genes transcribed from the same strand and with an intergenic distance of ≤50 bp were inferred to be co-transcribed based on developmental Cappable-seqencing (124) and RNA-sequencing (21, 23) data. Protein TMRs were predicted using DeepTMHMM (131). Protein functions were predicted using blastp (132, 133) and the AlphaFold Protein Structure Database (134, 135).

## ACKNOWLEDGMENTS

We thank Adam Deutschbauer for *E. coli* APA752.

This research was supported by National Science Foundation grant IOS-1951025 to L.K., by National Institutes of Health Award R35GM131762 to S.C., and by salary support for L.K. from Michigan State University AgBioResearch. A.F. was additionally supported by startup funds from the College of Natural Sciences and the School of Veterinary Medicine and AgBio Research at Michigan State University. The funders had no role in study design, data collection and interpretation, or the decision to submit the work for publication.

## REFERENCES

1. Kroos L, Wall D, Islam ST, Whitworth DE, Munoz-Dorado J, Higgs PI, Singer M, Mauriello EM, Treuner-Lange A, Sogaard-Andersen L, Kaimer C, Elias-Arnanz M, Stojkovic EA, Muller R, Volz C, Velicer GJ, Nan B. 2025. Milestones in the development of *Myxococcus xanthus* as a model multicellular bacterium. J Bacteriol 207:e0007125.

2. Kroos L. 2017. Highly signal-responsive gene regulatory network governing *Myxococcus* development. Trends Genet 33:3–15.

3. Dworkin M, Gibson SM. 1964. A system for studying microbial morphogenesis: rapid formation of microcysts in *Myxococcus xanthus*. Science 146:243–244.

4. Marcos-Torres FJ, Volz C, Muller R. 2020. An ambruticin-sensing complex modulates *Myxococcus xanthus* development and mediates myxobacterial interspecies communication. Nat Commun 11:5563.

5. Muller FD, Treuner-Lange A, Heider J, Huntley SM, Higgs PI. 2010. Global transcriptome analysis of spore formation in *Myxococcus xanthus* reveals a locus necessary for cell differentiation. BMC Genomics 11:264.

6. Wetmore KM, Price MN, Waters RJ, Lamson JS, He J, Hoover CA, Blow MJ, Bristow J, Butland G, Arkin AP, Deutschbauer A. 2015. Rapid quantification of mutant fitness in diverse bacteria by sequencing randomly bar-coded transposons. mBio 6:e00306–15.

7. Lee B, Holkenbrink C, Treuner-Lange A, Higgs PI. 2012. *Myxococcus xanthus* developmental cell fate production: heterogeneous accumulation of developmental regulatory proteins and reexamination of the role of MazF in developmental lysis. J Bacteriol 194:3058–3068.

8. Rajagopalan R, Kroos L. 2014. Nutrient-regulated proteolysis of MrpC halts expression of genes important for commitment to sporulation during *Myxococcus xanthus* development. J Bacteriol 196:2736–2747.

9. Saha S, Patra P, Igoshin O, Kroos L. 2019. Systematic analysis of the *Myxococcus xanthus* developmental gene regulatory network supports posttranslational regulation of FruA by C-signaling. Mol Microbiol 111:1732–1752.

10. Wireman JW, Dworkin M. 1975. Morphogenesis and developmental interactions in myxobacteria. Science 189:516–523.

11. Hodgkin J, Kaiser D. 1979. Genetics of gliding motility in *Myxococcus xanthus* (Myxobacterales): two gene systems control movement. Mol Gen Genet 171:177–191.

12. Zhang Z, Cotter CR, Lyu Z, Shimkets LJ, Igoshin OA. 2020. Data-driven models reveal mutant cell behaviors important for myxobacterial aggregation. mSystems 5:e00518–20.

13. Perez-Burgos M, Garcia-Romero I, Jung J, Schander E, Valvano MA, Sogaard-Andersen L. 2020. Characterization of the exopolysaccharide biosynthesis pathway in *Myxococcus xanthus*. J Bacteriol 202:e00335–20.

14. Rodriguez-Soto JP, Kaiser D. 1997. The *tgl* gene: social motility and stimulation in *Myxococcus xanthus*. J Bacteriol 179:4361–4371.

15. Nudleman E, Wall D, Kaiser D. 2005. Cell-to-cell transfer of bacterial outer membrane lipoproteins. Science 309:125–127.

16. Wall D. 2014. Molecular recognition in myxobacterial outer membrane exchange: functional, social and evolutionary implications. Mol Microbiol 91:209–220.

17. Siewering K, Jain S, Friedrich C, Webber-Birungi MT, Semchonok DA, Binzen I, Wagner A, Huntley S, Kahnt J, Klingl A, Boekema EJ, Sogaard-Andersen L, van der Does C. 2014. Peptidoglycan-binding protein TsaP functions in surface assembly of type IV pili. Proc Natl Acad Sci USA 111:E953–E961.

18. Chang YW, Rettberg LA, Treuner-Lange A, Iwasa J, Sogaard-Andersen L, Jensen GJ. 2016. Architecture of the type IVa pilus machine. Science 351:aad2001.

19. Treuner-Lange A, Chang YW, Glatter T, Herfurth M, Lindow S, Chreifi G, Jensen GJ, Sogaard-Andersen L. 2020. PilY1 and minor pilins form a complex priming the type IVa pilus in *Myxococcus xanthus*. Nat Commun 11:5054.

20. McLoon AL, Boeck ME, Bruckskotten M, Keyel AC, Sogaard-Andersen L. 2021. Transcriptomic analysis of the *Myxococcus xanthus* FruA regulon, and comparative developmental transcriptomic analysis of two fruiting body forming species, *Myxococcus xanthus* and *Myxococcus stipitatus*. BMC Genomics 22:784.

21. Munoz-Dorado J, Moraleda-Munoz A, Marcos-Torres FJ, Contreras-Moreno FJ, Martin-Cuadrado AB, Schrader JM, Higgs PI, Perez J. 2019. Transcriptome dynamics of the *Myxococcus xanthus* multicellular developmental program. eLife 8:e50374.

22. Mahanta U, Wassmuth R, Brighty S, Treuner-Lange A, Sharma G. 2026. Diversity, classification, and evolution of myxobacterial PilY1 proteins. Front Microbiol 17:1826482.

23. Farrugia MA, Rajagopalan R, Kroos L. 2025. Transcriptomic analysis of *Myxococcus xanthus csgA, fruA, and mrpC* mutants reveals extensive and diverse roles of key regulators in the multicellular developmental process. BMC Genomics 26:355.

24. Luciano J, Agrebi R, Le Gall AV, Wartel M, Fiegna F, Ducret A, Brochier-Armanet C, Mignot T. 2011. Emergence and modular evolution of a novel motility machinery in bacteria. PLoS Genet 7:e1002268.

25. Wartel M, Ducret A, Thutupalli S, Czerwinski F, Le Gall AV, Mauriello EM, Bergam P, Brun YV, Shaevitz J, Mignot T. 2013. A versatile class of cell surface directional motors gives rise to gliding motility and sporulation in *Myxococcus xanthus*. PLoS Biol 11:e1001728.

26. Muller FD, Schink CW, Hoiczyk E, Cserti E, Higgs PI. 2012. Spore formation in *Myxococcus xanthus* is tied to cytoskeleton functions and polysaccharide spore coat deposition. Mol Microbiol 83:486–505.

27. Pathak DT, Wall D. 2012. Identification of the *cglC, cglD, cglE*, and *cglF* genes and their role in cell contact-dependent gliding motility in *Myxococcus xanthus*. J Bacteriol 194:1940–1949.

28. Hodgkin J, Kaiser D. 1977. Cell-to-cell stimulation of motility in nonmotile mutants of *Myxococcus*. Proc Natl Acad Sci USA 74:2938–2942.

29. Pathak DT, Wei X, Bucuvalas A, Haft DH, Gerloff DL, Wall D. 2012. Cell contact-dependent outer membrane exchange in myxobacteria: genetic determinants and mechanism. PLoS Genet 8:e1002626.

30. Cambillau C, Mignot T. 2025. Structural model of a bacterial focal adhesion complex. Commun Biol 8:119.

31. White DJ, Hartzell PL. 2000. AglU, a protein required for gliding motility and spore maturation of *Myxococcus xanthus*, is related to WD-repeat proteins. Mol Microbiol 36:662–678.

32. Ramirez Carbo CA, Faromiki OG, Nan B. 2024. A lytic transglycosylase connects bacterial focal adhesion complexes to the peptidoglycan cell wall. eLife 13:RP99273.

33. Ramirez Carbo CA, Irazoki O, Venkatesan S, Chen LJS, Morales HA, Garcia Avila AJ, Cheung HL, Cava F, Nan B. 2025. A novel mechanism for bacterial sporulation based on programmed peptidoglycan degradation. eLife 14:RP108250.

34. Leonardy S, Miertzschke M, Bulyha I, Sperling E, Wittinghofer A, Sogaard-Andersen L. 2010. Regulation of dynamic polarity switching in bacteria by a Ras-like G-protein and its cognate GAP. EMBO J 29:2276–2289.

35. Zhang Y, Franco M, Ducret A, Mignot T. 2010. A bacterial Ras-like small GTP-binding protein and its cognate GAP establish a dynamic spatial polarity axis to control directed motility. PLoS Biol 8:e1000430.

36. Oklitschek M, Carreira LAM, Muratoglu M, Sogaard-Andersen L, Treuner-Lange A. 2024. Combinatorial control of type IVa pili formation by the four polarized regulators MglA, SgmX, FrzS, and SopA. J Bacteriol 206:e0010824.

37. Mercier R, Bautista S, Delannoy M, Gibert M, Guiseppi A, Herrou J, Mauriello EMF, Mignot T. 2020. The polar Ras-like GTPase MglA activates type IV pilus via SgmX to enable twitching motility in *Myxococcus xanthus*. Proc Natl Acad Sci USA 117:28366–28373.

38. Potapova A, Carreira LAM, Sogaard-Andersen L. 2020. The small GTPase MglA together with the TPR domain protein SgmX stimulates type IV pili formation in *M. xanthus*. Proc Natl Acad Sci USA 117:23859–23868.

39. Hartzell P, Kaiser D. 1991. Upstream gene of the *mgl* operon controls the level of MglA protein in *Myxococcus xanthus*. J Bacteriol 173:7625–7635.

40. Zusman DR. 1982. "Frizzy" mutants: a new class of aggregation-defective developmental mutants of *Myxococcus xanthus*. J Bacteriol 150:1430–1437.

41. Blackhart BD, Zusman DR. 1985. "Frizzy" genes of *Myxococcus xanthus* are involved in control of frequency of reversal of gliding motility. Proc Natl Acad Sci USA 82:8767–8770.

42. Bustamante VH, Martinez-Flores I, Vlamakis HC, Zusman DR. 2004. Analysis of the Frz signal transduction system of *Myxococcus xanthus* shows the importance of the conserved C-terminal region of the cytoplasmic chemoreceptor FrzCD in sensing signals. Mol Microbiol 53:1501–1513.

43. Kashefi K, Hartzell P. 1995. Genetic supression and phenotypic masking of a *Myxococcus xanthus frzF*^-^ defect. Mol Microbiol 15:483–494.

44. Hoang Y, Franklin JL, Dufour YS, Kroos L. 2021. Cell density, alignment, and orientation correlate with C-signal-dependent gene expression during *Myxococcus xanthus* development. Proc Natl Acad Sci USA 118:e2111706118.

45. Kearns DB, Venot A, Bonner PJ, Stevens B, Boons GJ, Shimkets LJ. 2001. Identification of a developmental chemoattractant in *Myxococcus xanthus* through metabolic engineering. Proc Natl Acad Sci USA 98:13990–13994.

46. Bhat S, Ahrendt T, Dauth C, Bode HB, Shimkets LJ. 2014. Two lipid signals guide fruiting body development of *Myxococcus xanthus*. mBio 5:e00939–13.

47. Lorenzen W, Ahrendt T, Bozhuyuk KA, Bode HB. 2014. A multifunctional enzyme is involved in bacterial ether lipid biosynthesis. Nat Chem Biol 10:425–427.

48. Boynton TO, Shimkets LJ. 2015. *Myxococcus* CsgA, Drosophila Sniffer and human HSD17B10 are cardiolipin phospholipases. Genes Dev 29:1903-1914.

49. Gill RE, Karlok M, Benton D. 1993. *Myxococcus xanthus* encodes an ATP-dependent protease which is required for developmental gene transcription and intercellular signaling. J Bacteriol 175:4538–4544.

50. Tojo N, Inouye S, Komano T. 1993. The *lonD* gene is homologous to the *lon* gene encoding an ATP-dependent protease and is essential for the development of *Myxococcus xanthus*. J Bacteriol 175:4545–4549.

51. Gill RE, Cull MG. 1986. Control of developmental gene expression by cell-to-cell interactions in *Myxococcus xanthus*. J Bacteriol 168:341–347.

52. Kroos L, Kaiser D. 1987. Expression of many developmentally regulated genes in *Myxococcus* depends on a sequence of cell interactions. Genes Dev 1:840–854.

53. Bhat S, Boynton TO, Pham D, Shimkets LJ. 2014. Fatty acids from membrane lipids become incorporated into lipid bodies during *Myxococcus xanthus* differentiation. PLoS One 9:e99622.

54. Hoang Y, Franklin J, Dufour YS, Kroos L. 2024. Short-range C-signaling restricts cheating behavior during *Myxococcus xanthus* development. mBio 15:e0244024.

55. Nielsen M, Rasmussen AA, Ellehauge E, Treuner-Lange A, Sogaard-Andersen L. 2004. HthA, a putative DNA-binding protein, and HthB are important for fruiting body morphogenesis in *Myxococcus xanthus*. Microbiology 150:2171–2183.

56. Perez-Burgos M, Sogaard-Andersen L. 2020. Biosynthesis and function of cell-surface polysaccharides in the social bacterium *Myxococcus xanthus*. Biol Chem 401:1375–1387.

57. Yang Z, Ma X, Tong L, Kaplan HB, Shimkets LJ, Shi W. 2000. *Myxococcus xanthus dif* genes are required for biogenesis of cell surface fibrils essential for social gliding motility. J Bacteriol 182:5793–5798.

58. Black WP, Xu Q, Yang Z. 2006. Type IV pili function upstream of the Dif chemotaxis pathway in *Myxococcus xanthus* EPS regulation. Mol Microbiol 61:447–456.

59. Li Y, Sun H, Ma X, Lu A, Lux R, Zusman D, Shi W. 2003. Extracellular polysaccharides mediate pilus retraction during social motility of *Myxococcus xanthus*. Proc Natl Acad Sci USA 100:5443–5448.

60. Bellenger K, Ma X, Shi W, Yang Z. 2002. A CheW homologue is required for *Myxococcus xanthus* fruiting body development, social gliding motility, and fibril biogenesis. J Bacteriol 184:5654–5660.

61. Yang Z, Geng Y, Xu D, Kaplan HB, Shi W. 1998. A new set of chemotaxis homologues is essential for *Myxococcus xanthus* social motility. Mol Microbiol 30:1123–1130.

62. Shimkets LJ. 1986. Role of cell cohesion in *Myxococcus xanthus* fruiting body formation. J Bacteriol 166:842–848.

63. Chang BY, Dworkin M. 1994. Isolated fibrils rescue cohesion and development in the Dsp mutant of *Myxococcus xanthus*. J Bacteriol 176:7190–7196.

64. Black WP, Yang Z. 2004. *Myxococcus xanthus* chemotaxis homologs DifD and DifG negatively regulate fibril polysaccharide production. J Bacteriol 186:1001–1008.

65. Yang Z, Geng Y, Shi W. 1998. A DnaK homolog in *Myxococcus xanthus* is involved in social motility and fruiting body formation. J Bacteriol 180:218–224.

66. Weimer RM, Creighton C, Stassinopoulos A, Youderian P, Hartzell PL. 1998. A chaperone in the HSP70 family controls production of extracellular fibrils in *Myxococcus xanthus*. J Bacteriol 180:5357–5368.

67. Pan Z, Zhuo L, Wan Ty, Chen Ry, Li Yz. 2024. DnaK duplication and specialization in bacteria correlates with increased proteome complexity. mSystems 9:e0115423.

68. Pan Z, Zhang Z, Zhuo L, Wan TY, Li YZ. 2021. Bioinformatic and functional characterization of Hsp70s in *Myxococcus xanthus*. mSphere 6:e00305–21.

69. Molinari G, Ribeiro SS, Muller K, Mayer BE, Rohde M, Arce-Rodriguez A, Vargas-Guerrero JJ, Avetisyan A, Wissing J, Tegge W, Jansch L, Bronstrup M, Danchin A, Jahn M, Timmis KN, Ebbinghaus S, Jahn D, Borrero-de Acuna JM. 2025. Multiple chaperone DnaK-FliC flagellin interactions are required for *Pseudomonas aeruginosa* flagellum assembly and indicate a new function for DnaK. Microb Biotechnol 18:e70096.

70. McDonald HJ, Kweon H, Kurnfuli S, Risser DD. 2022. A DnaK(Hsp70) chaperone system connects type IV pilus activity to polysaccharide secretion in cyanobacteria. mBio 13:e0051422.

71. Youderian P, Hartzell PL. 2006. Transposon insertions of *magellan-4* that impair social gliding motility in *Myxococcus xanthus*. Genetics 172:1397–1410.

72. Pham VD, Shebelut CW, Mukherjee B, Singer M. 2005. RasA is required for *Myxococcus xanthus* development and social motility. J Bacteriol 187:6845–6848.

73. Xue S, Mercier R, Guiseppi A, Kosta A, De Cegli R, Gagnot S, Mignot T, Mauriello EMF. 2022. The differential expression of PilY1 proteins by the HsfBA phosphorelay allows twitching motility in the absence of exopolysaccharides. PLoS Genet 18:e1010188.

74. Ueki T, Inouye S. 2002. Transcriptional activation of a heat-shock gene, *lonD*, of *Myxococcus xanthus* by a two component histidine-aspartate phosphorelay system. J Biol Chem 277:6170–6177.

75. Volz C, Kegler C, Muller R. 2012. Enhancer binding proteins act as hetero-oligomers and link secondary metabolite production to myxococcal development, motility, and predation. Chem Biol 19:1447–1459.

76. Caberoy NB, Welch RD, Jakobsen JS, Slater SC, Garza AG. 2003. Global mutational analysis of NtrC-like activators in *Myxococcus xanthus*: identifying activator mutants defective for motility and fruiting body development. J Bacteriol 185:6083–6094.

77. Moak PL, Black WP, Wallace RA, Li Z, Yang Z. 2015. The Hsp70-like StkA functions between T4P and Dif signaling proteins as a negative regulator of exopolysaccharide in *Myxococcus xanthus*. PeerJ 3:e747.

78. Yu R, Kaiser D. 2007. Gliding motility and polarized slime secretion. Mol Microbiol 63:454–467.

79. Perez-Burgos M, Garcia-Romero I, Jung J, Valvano MA, Sogaard-Andersen L. 2019. Identification of the lipopolysaccharide O-antigen biosynthesis priming enzyme and the O-antigen ligase in *Myxococcus xanthus*: critical role of LPS O-antigen in motility and development. Mol Microbiol 112:1178–1198.

80. Gong Y, Zhang Z, Zhou XW, Anwar MN, Hu XZ, Li ZS, Chen XJ, Li YZ. 2018. Competitive interactions between incompatible mutants of the social bacterium *Myxococcus xanthus* DK1622. Front Microbiol 9:1200.

81. Holkenbrink C, Hoiczyk E, Kahnt J, Higgs PI. 2014. Synthesis and assembly of a novel glycan layer in *Myxococcus xanthus* spores. J Biol Chem 289:32364-32378.

82. Perez-Burgos M, Garcia-Romero I, Valvano MA, Sogaard Andersen L. 2020. Identification of the Wzx flippase, Wzy polymerase and sugar-modifying enzymes for spore coat polysaccharide biosynthesis in *Myxococcus xanthus*. Mol Microbiol 113:1189–1208.

83. Giglio KM, Zhu C, Klunder C, Kummer S, Garza AG. 2015. The enhancer binding protein Nla6 regulates developmental genes that are important for *Myxococcus xanthus* sporulation. J Bacteriol 197:1276–1287.

84. Saha S, Kroos L. 2024. Regulation of late-acting operons by three transcription factors and a CRISPR-Cas component during *Myxococcus xanthus* development. Mol Microbiol 121:1002–1020.

85. Fink JM, Zissler JF. 1989. Defects in motility and development of *Myxococcus xanthus* lipopolysaccharide mutants. J Bacteriol 171:2042–2048.

86. Vassallo C, Pathak DT, Cao P, Zuckerman DM, Hoiczyk E, Wall D. 2015. Cell rejuvenation and social behaviors promoted by LPS exchange in myxobacteria. Proc Natl Acad Sci USA 112:E2939–46.

87. Nariya H, Inouye M. 2008. MazF, an mRNA interferase, mediates programmed cell death during multicellular *Myxococcus* development. Cell 132:55–66.

88. Boynton TO, McMurry JL, Shimkets LJ. 2013. Characterization of *Myxococcus xanthus* MazF and implications for a new point of regulation. Mol Microbiol 87:1267–1276.

89. Kaimer C, Weltzer ML, Wall D. 2023. Two reasons to kill: predation and kin discrimination in myxobacteria. Microbiology (Reading) 169:001372.

90. Wang F, Luo J, Zhang Z, Liu Y, Sheng D, Zhuo L, Li Yz. 2025. Differential crosstalk between toxin-immunity protein homologs divides *Myxococcus* nonself siblings into close and distant social relatives. mBio 16:e0390224.

91. Gong Y, Zhang Z, Liu Y, Zhou XW, Anwar MN, Li ZS, Hu W, Li YZ. 2018. A nuclease-toxin and immunity system for kin discrimination in *Myxococcus xanthus*. Environ Microbiol 20:2552–2567.

92. Vassallo CN, Wall D. 2019. Self-identity barcodes encoded by six expansive polymorphic toxin families discriminate kin in myxobacteria. Proc Natl Acad Sci USA 116:24808–24818.

93. Troselj V, Treuner-Lange A, Sogaard-Andersen L, Wall D. 2018. Physiological heterogeneity triggers sibling conflict mediated by the type VI secretion system in an aggregative multicellular bacterium. mBio 9:e01645–17.

94. Vassallo CN, Troselj V, Weltzer ML, Wall D. 2020. Rapid diversification of wild social groups driven by toxin-immunity loci on mobile genetic elements. ISME J 14:2474–2487.

95. Wielgoss S, Didelot X, Chaudhuri RR, Liu X, Weedall GD, Velicer GJ, Vos M. 2016. A barrier to homologous recombination between sympatric strains of the cooperative soil bacterium *Myxococcus xanthus*. ISME J 10:2468–2477.

96. Sah GP, Cao P, Wall D. 2020. MYXO-CTERM sorting tag directs proteins to the cell surface via the type II secretion system. Mol Microbiol 113:1038–1051.

97. Velicer GJ, Kroos L, Lenski RE. 2000. Developmental cheating in the social bacterium *Myxococcus xanthus*. Nature 404:598–601.

98. Fiegna F, Velicer GJ. 2005. Exploitative and hierarchical antagonism in a cooperative bacterium. PLoS Biol 3:e370.

99. Fiegna F, Yu YT, Kadam SV, Velicer GJ. 2006. Evolution of an obligate social cheater to a superior cooperator. Nature 441:310–314.

100. Yu YT, Yuan X, Velicer GJ. 2010. Adaptive evolution of an sRNA that controls *Myxococcus* development. Science 328:993.

101. La Fortezza M, Verwilt J, Cossey SM, Eisner SA, Velicer GJ, Yu YN. 2025. Deletion of an sRNA primes development in a multicellular bacterium. iScience 28:111980.

102. Laue BE, Gill RE. 1994. Use of a phase variation-specific promoter of *Myxococcus xanthus* in a strategy for isolating a phase-locked mutant. J Bacteriol 176:5341–5349.

103. Laue BE, Gill RE. 1995. Using a phase-locked mutant of *Myxococcus xanthus* to study the role of phase variation in development. J Bacteriol 177:4089–4096.

104. Dziewanowska K, Settles M, Hunter S, Linquist I, Schilkey F, Hartzell PL. 2014. Phase variation in *Myxococcus xanthus* yields cells specialized for iron sequestration. PLoS One 9:e95189.

105. Meiser P, Bode HB, Muller R. 2006. The unique DKxanthene secondary metabolite family from the myxobacterium *Myxococcus xanthus* is required for developmental sporulation. Proc Natl Acad Sci USA 103:19128–19133.

106. Kuspa A, Kaiser D. 1989. Genes required for developmental signaling in *Myxococcus xanthus*: three *asg* loci. J Bacteriol 171:2762–2772.

107. Overgaard M, Wegener-Feldbrugge S, Sogaard-Andersen L. 2006. The orphan response regulator DigR is required for synthesis of extracellular matrix fibrils in *Myxococcus xanthus*. J Bacteriol 188:4384–4394.

108. Nicolas FJ, Ruiz-Vazquez RM, Murillo FJ. 1994. A genetic link between light response and multicellular development in the bacterium *Myxococcus xanthus*. Genes Dev 8:2375–2387.

109. Plamann L, Davis J, Cantwell B, Mayor J. 1994. Evidence that *asgB* encodes a DNA-binding protein essential for growth and development of *Myxococcus xanthus*. J Bacteriol 176:2013–2022.

110. Li Y, Plamann L. 1996. Purification and in vitro phosphorylation of *Myxococcus xanthus* AsgA protein. J Bacteriol 178:289–292.

111. Konovalova A, Wegener-Feldbrugge S, Sogaard-Andersen L. 2012. Two intercellular signals required for fruiting body formation in *Myxococcus xanthus* act sequentially but non-hierarchically. Mol Microbiol 86:65–81.

112. Kuspa A, Plamann L, Kaiser D. 1992. A-signalling and the cell density requirement for *Myxococcus xanthus* development. J Bacteriol 174:7360–7369.

113. Bernal-Bernal D, Abellon-Ruiz J, Iniesta AA, Pajares-Martinez E, Bastida-Martinez E, Fontes M, Padmanabhan S, Elias-Arnanz M. 2018. Multifactorial control of the expression of a CRISPR-Cas system by an extracytoplasmic function σ/anti-σ pair and a global regulatory complex. Nucleic Acids Res 46:6726–6745.

114. Furusawa G, Dziewanowska K, Stone H, Settles M, Hartzell P. 2011. Global analysis of phase variation in *Myxococcus xanthus*. Mol Microbiol 81:784–804.

115. Moine A, Agrebi R, Espinosa L, Kirby JR, Zusman DR, Mignot T, Mauriello EM. 2014. Functional organization of a multimodular bacterial chemosensory apparatus. PLoS Genet 10:e1004164.

116. Xiao Y, Wall D. 2014. Genetic redundancy, proximity, and functionality of *lspA*, the target of antibiotic TA, in the *Myxococcus xanthus* producer strain. J Bacteriol 196:1174–1183.

117. Rolbetzki A, Ammon M, Jakovljevic V, Konovalova A, Sogaard-Andersen L. 2008. Regulated secretion of a protease activates intercellular signaling during fruiting body formation in *M. xanthus*. Dev Cell 15:627–634.

118. Darnell CL, Wilson JM, Tiwari N, Fuentes EJ, Kirby JR. 2014. Chemosensory regulation of a HEAT-repeat protein couples aggregation and sporulation in *Myxococcus xanthus*. J Bacteriol 196:3160–3168.

119. Chen IK, Satinsky BM, Velicer GJ, Yu YN. 2019. sRNA-pathway genes regulating myxobacterial development exhibit clade-specific evolution. Evol Dev 21:82–95.

120. Nair RR, Fiegna F, Velicer GJ. 2018. Indirect evolution of social fitness inequalities and facultative social exploitation. Proc R Soc B 285:20180054.

121. Dey A, Wall D. 2014. A genetic screen in *Myxococcus xanthus* identifies mutants that uncouple outer membrane exchange from a downstream cellular response. J Bacteriol 196:4324–4332.

122. Wright TA, Jiang L, Park JJ, Anderson WA, Chen G, Hallberg ZF, Nan B, Hammond MC. 2020. Second messengers and divergent HD-GYP phosphodiesterases regulate 3’,3’-cGAMP signaling. Mol Microbiol 113:222–236.

123. Skotnicka D, Smaldone GT, Petters T, Trampari E, Liang J, Kaever V, Malone JG, Singer M, Sogaard-Andersen L. 2016. A minimal threshold of c-di-GMP is essential for fruiting body formation and sporulation in *Myxococcus xanthus*. PLoS Genet 12:e1006080.

124. Kuzmich S, Blumenkamp P, Meier D, Szadkowski D, Goesmann A, Becker A, Sogaard-Andersen L. 2022. CRP-Like transcriptional regulator MrpC curbs c-di-GMP and 3’,3’-cGAMP nucleotide levels during development in *Myxococcus xanthus*. mBio 13:e00044–22.

125. Liu Y, Wang J, Zhang Z, Wang F, Gong Y, Sheng DH, Li YZ. 2021. Two PAAR proteins with different C-terminal extended domains have distinct ecological functions in *Myxococcus xanthus*. Appl Environ Microbiol 87:e00080–21.

126. Viswanathan P, Murphy K, Julien B, Garza AG, Kroos L. 2007. Regulation of *dev*, an operon that includes genes essential for *Myxococcus xanthus* development and CRISPR-associated genes and repeats. J Bacteriol 189:3738–3750.

127. Rajagopalan R, Kroos L. 2017. The *dev* operon regulates the timing of sporulation during *Myxococcus xanthus* development. J Bacteriol 199:e00788–16.

128. Rajagopalan R, Wielgoss S, Lippert G, Velicer GJ, Kroos L. 2015. *devI* is an evolutionarily young negative regulator of *Myxococcus xanthus* development. J Bacteriol 197:1249–1262.

129. Kaiser D. 1979. Social gliding is correlated with the presence of pili in *Myxococcus xanthus*. Proc Natl Acad Sci USA 76:5952–5956.

130. Fiebig A, Schnizlein MK, Pena-Rivera S, Trigodet F, Dubey AA, Hennessy MK, Basu A, Pott S, Dalal S, Rubin D, Sogin ML, Eren AM, Chang EB, Crosson S. 2024. Bile acid fitness determinants of a *Bacteroides fragilis* isolate from a human pouchitis patient. mBio 15:e0283023.

131. Hallgren J, Tsirigos K, Pedersen M, Armenteros J, Marcatili P, Nielsen H, Krogh A, Winther O. 2022. DeepTMHMM predicts alpha and beta transmembrane proteins using deep neural networks. bioRxiv 10.1101/2022.04.08.487609

132. Altschul SF, Gish W, Miller W, Myers EW, Lipman DJ. 1990. Basic local alignment search tool. J Mol Biol 215:403–410.

133. blastp. 2026. Protein [Internet]. Bethesda (MD): National Library of Medicine (US), National Center for Biotechnology Information; [1988] - [cited 2026 July 15] Available from: https://wwwncbinlmnihgov/protein/.

134. Jumper J, Evans R, Pritzel A, Green T, Figurnov M, Ronneberger O, Tunyasuvunakool K, Bates R, Zidek A, Potapenko A, Bridgland A, Meyer C, Kohl SAA, Ballard AJ, Cowie A, Romera-Paredes B, Nikolov S, Jain R, Adler J, Back T, Petersen S, Reiman D, Clancy E, Zielinski M, Steinegger M, Pacholska M, Berghammer T, Bodenstein S, Silver D, Vinyals O, Senior AW, Kavukcuoglu K, Kohli P, Hassabis D. 2021. Highly accurate protein structure prediction with AlphaFold. Nature 596:583–589.

135. Varadi M, Bertoni D, Magana P, Paramval U, Pidruchna I, Radhakrishnan M, Tsenkov M, Nair S, Mirdita M, Yeo J, Kovalevskiy O, Tunyasuvunakool K, Laydon A, Zidek A, Tomlinson H, Hariharan D, Abrahamson J, Green T, Jumper J, Birney E, Steinegger M, Hassabis D, Velankar S. 2024. AlphaFold Protein Structure Database in 2024: providing structure coverage for over 214 million protein sequences. Nucleic Acids Res 52:D368–D375.

