## Supporting Information for "Temporal, genome-scale analysis of *Myxococcus xanthus* developmental fate in a mixed population"

Running title: Barcoded mutant analysis of *M. xanthus* development

**Results and Discussion**

**Genes involved in the regulation of motility reversal**

In previous work, a *pglH* mutant was partially defective in gliding motility and reversed its direction of movement at twice the normal frequency, leading Yu and Kaiser (1) to propose that PglH is a component of a reversal clock. If so, a *pglH* mutant is expected to be defective in T4P-dependent motility as well as gliding motility. Insertions in *pglH* (MXAN_2050) had a fitness profile resembling that of *frzF/E/B* (Fig. 2B), indicating aggregation and sporulation defects consistent with a role in regulating motility reversal. PglH is an orphan response regulator (RR) (i.e., no histidine kinase [HK] is encoded nearby) with a tetratricopeptide repeat (TPR) domain, suggesting a role in assembly of a protein complex. A *plpA* mutant also reversed motility at twice the normal frequency, and was defective in both types of motility and localization of MglA (2). Despite these defects, the *plpA* mutant formed aggregates on starvation agar, although the number of aggregates appeared to be reduced and sporulation was not measured. Insertions in *plpA* (MXAN_2528) produced a fitness pattern indicating aggregation and sporulation defects in the mixed population under submerged culture (Fig. 2B), consistent with a role in regulating motility reversal. Although PlpA is a PilZ-like protein, its regulatory function does not require cyclic-di-GMP (c-di-GMP), and PlpA was shown to interact directly with AglS (2).

We did not expect *frgA* insertions to cause developmental defects in the mixed population because the frizzy phenotype of a *frgA* mutant was extracellularly complemented for aggregation and sporulation, suggesting a defect in production of an extracellular signal supplied by wild type upon co-development (3). However, insertions in *frgA* (MXAN_1130) showed fitness effects similar to insertions in the frizzy genes *frzF/E/B* (Fig. 2B), indicating that aggregation and sporulation were not extracellularly complemented in our mutant pool, and suggesting instead that FrgA cell-autonomously regulates motility reversal. FrgA is predicted to be a 942-residue soluble protein with an N-terminal GspE/PulE/PilB domain. The function of FrgA in control of motility reversal remains to be determined.

**Signal-dependent regulators and uncharacterized genes**

PilT is the ATPase for T4P retraction (4). Strains with insertions in *pilT* (MXAN_5787) were enriched in agg samples at 12-30 h (earlier than other *pil* mutants), accumulated in non-agg samples less than other *pil* mutants, and exhibited variable sporulation defects: slight overrepresentation in time course 2, but underrepresentation similar to other T4P mutants in time course 3 (Fig. 2B). Our results provide evidence that T4P extension without retraction allows greater aggregation of *M. xanthus* in mixed populations at early stages of development, consistent with the observation that a *pilT* mutant aggregates better than *pilH* and *pilA* mutants (5). Insertions in *sglT* (MXAN_3284) had similar fitness effects as insertions in *pilT* (Fig. 2B). Like PilT, SglT appears to function in T4P retraction, but SglT is dispensable at lower growth temperatures (6). SglT is annotated as an $\alpha/\beta$ hydrolase, but its enzymatic activity is unknown. Development of an *sglT* mutant has not been reported.

Two sets of two-component signaling genes lie in the T4P locus, the sensor HK genes *pilS* and *pilS2*, and the RR genes *pilR* and *pilR2*. The fitness profiles of *pilR* (MXAN_5784) and *pilR2* (MXAN_5777) insertions cluster with other *pil* genes (Fig. 2B), consistent with the requirement of both RRs for T4P-dependent motility (7-9). PilR is required for *pilA* expression (8) and PilR2 positively regulates T4P assembly and exopolysaccharide (EPS) production (7), which in turn stimulates T4P retraction (10). PilS negatively regulates *pilA* expression (8), suggesting that it acts as a phosphatase of PilR. Insertions in *pilS* (MXAN_5785) did not exhibit significant defects in our experiments (Table S2). PilS2 specifically phosphorylates PilR2 *in vitro* (11), but in-frame deletion of *pilS2* had no effect on T4P-dependent motility during growth (7). The role of *pilS2* in development was not tested previously. Insertions in *pilS2* (MXAN_5778) gave fitness patterns resembling those of *pilT* and *sglT*, but with dampened magnitude (Fig. 2B), consistent with a role in regulating T4P assembly and/or retraction.

Insertions in *sdeK* (MXAN_1014) and *fruA* (MXAN_3117) caused fitness patterns similar to insertions in *pilT* and *frzS*, including little or no effect on glycerol-induced sporulation (Fig. 2B). SdeK is an orphan HK and FruA is an orphan RR whose partners are unknown. Previous work showed that *sdeK* and *fruA* mutants are severely impaired in starvation-induced aggregation, sporulation, and gene expression (12-17).

Insertions in the *pktA5* (MXAN_0930), *espB* (MXAN_0932), and *pktB8* (MXAN_0933) gene cluster yielded fitness profiles indicating aggregation defects, variable starvation-induced sporulation, and weakly impaired glycerol-induced sporulation (Fig. 2B). Defined mutants had impaired aggregation and sporulation in a density-dependent fashion (18). A mutation in *espB* delayed aggregation and sporulation, whereas an *espA* (MXAN_0931) mutation accelerated development (19). EspB is a putative oligopeptide transporter, and PktA5 and PktB8 are serine/threonine protein kinases (STPKs) that interact with EspA, a hybrid HK-RR, in a signal transduction pathway that regulates developmental progression (18-21). We did not obtain fitness scores for *espA*.

Our dataset revealed five largely uncharacterized genes whose insertions produced fitness patterns consistent with reduced aggregation and sporulation (Fig. 2B). One gene encodes a signal transduction protein; MXAN_0399 is an HK that is not transcriptionally regulated during the first 24 h of development (22). The other four genes encode predicted Gcn5-related N-acetyltransferase (GNAT) (MXAN_3054), metallophosphoesterase (MXAN_2613), amino acid dehydrogenase (MXAN_5873), and aldehyde dehydrogenase (MXAN_0625) enzymes. Creating in-frame deletion mutants and examining their development in monoculture and in mixtures with wild type would likely provide functional insights into these previously unstudied genes.

**Genes impacting cell loss during development**

MXAN_0234 encodes a putative inner membrane (IM) anti-$\sigma$ factor with a C-terminal cytoplasmic domain that likely inhibits extracytoplasmic function $\sigma$ factor MXAN_0233 (23), encoded upstream and likely co-transcribed, but its regulon is uncharacterized. Both proteins are conserved in the Myxococcales. Strains with insertions in MXAN_0234 were depleted in developmental samples (Fig. 4B). Insertions in MXAN_0233 were not sufficiently represented in our pool. MXAN_3963 encodes a protein similar to glycine/D-amino acid oxidase DadA. Upstream and likely co-transcribed is MXAN_3962, encoding an iron-sulfur cluster-binding protein. The two proteins are conserved in the Myxococcales and may function together, but are uncharacterized. Strains with insertions in either gene were similarly depleted in starved samples (Fig. 4B), indicating modest defects in attachment and/or survival. MXAN_3598 encodes a predicted DNA cytosine methyltransferase located downstream of MXAN_3599, encoding a putative very short patch repair endonuclease. Insertions in MXAN_3599 did not exhibit defects (Table S2), but insertions in MXAN_3598 were underrepresented in both starvation- and glycerol-induced samples (Fig. 4B). These two genes have not been studied in *M. xanthus*, but the predicted enzymes typically repair T/G mismatches resulting from 5-methylcytosine deamination to thymine in bacteria. In *E. coli*, this very short patch repair process is efficient during stationary phase (24) and aids genome stability (25). Our results suggest that DNA repair contributes to attachment and/or survival during *M. xanthus* development.

**Genes in the second group of developmental winners**

PopC is the protease that cleaves the C-signal precursor CsgA/p25 to its active p17 form extracellularly during starvation (26). We did not expect *popC* mutants to form spores in our experiments, because a *popC* mutant failed to form spores alone or upon co-development with wild type at an initial ratio of 1:1, suggesting a cell-autonomous requirement for PopC in addition to its extracellular role in C-signal production (26). However, strains with insertions in *popC* (MXAN_0206) were enriched in starvation-induced spore samples (Fig. 5A), indicating that *popC* mutants cheated in the presence of excess *popC*$^+$ cells in our Tn-Himar mutant pool. *csgA* mutants have been reported to cheat in some experiments, but the outcome depends on the mixing ratio with wild type (27, 28) and other factors (29). In our experiments, insertions in *csgA* (MXAN_1294) did not exhibit defects (Table S2).

The anti-$\sigma$ factor DdvA regulates DdvS, a $\sigma$ factor primarily involved in activating a CRISPR-Cas system, and a *ddvA* mutant forms about half as many fruiting bodies as wild type (30). Our finding that insertions in *ddvA* (MXAN_7288) were consistently overrepresented in spore samples (Fig. 5A) is therefore surprising and warrants further investigation. Insertions in *ddvS* (MXAN_7289) did not exhibit developmental defects in our experiments (Table S2).

CheB7 is a predicted methylesterase whose loss presumably enhances signaling through the Che7 chemosensory system implicated in coupling aggregation and sporulation (31). In-frame deletion of *cheB7* delayed aggregation, but the sporulation level was normal. Insertions in *cheB7* (MXAN_6959) did not impact fitness scores in non-agg and agg samples, but led to enrichment in starvation-induced spore samples (Fig. 5A).

*Nla7*/MXAN_0938 encode an RR/HK pair that is not transcriptionally regulated during the first 24 h of development (22). An *nla7* mutant exhibits normal aggregation and sporulation in monoculture (9), but insertions in *nla7* (MXAN_0937) or MXAN_0938 were enriched in spore samples (Fig. 5A). Altogether, the results suggest that the spore enrichment we observed was due to social interactions during development of the Tn-Himar mutant pool. A defined MXAN_0938 mutant has not been reported.

The second group of developmental winners includes 17 other genes without reported defined mutants. MXAN_5715 is transcriptionally induced early in development and encodes an orphan hybrid RR-HK-like protein (22) that is otherwise uncharacterized. Insertions in MXAN_5715 were overrepresented ~8-fold in starvation-induced spores and >2-fold in glycerol-induced spores (Fig. 5A). Consistent with the extensive surface remodeling required for differentiation into resistant spores, insertion strains disrupted in predicted lipoproteins (MXAN_6899, 1965) were enriched in spore samples (Fig. 5A), as was a glycosyltransferase (MXAN_1036) encoded in the BPS gene cluster (32). Strains with insertions in 13 other genes were also enriched in spore samples (Fig. 5A). These genes encode enzymes with predicted functions in protein disulfide bond formation (MXAN_2266), short-chain molecule oxidation/reduction (MXAN_1947, 3020), nucleic acid metabolism (MXAN_1053, 5871, 6077), ribosomal protein S6 glutamalyation (MXAN_0352, 3061), and small cytosolic peptide degradation (MXAN_2016), as well as a putative sodium-coupled membrane transporter (MXAN_4237), a putative molybdenum cofactor carrier protein (MXAN_4547), and proteins lacking significant similarity to proteins of known function (MXAN_0330, 2784).

**Genes in the third group of developmental winners**

LtgA is a lytic transglycosylase necessary for PG degradation during glycerol-induced sporulation (33). An *ltgA* mutant aggregated normally and starvation-induced spores were observed but not quantified. In future studies, spore quantification of an *ltgA* mutant in monoculture could clarify why insertions in *ltgA* were overrepresented in our starvation-induced spore samples (Fig. 5A).

To our knowledge, a defined MXAN_2985 mutant has not been reported. MXAN_2985 encodes a predicted Fic family protein, likely involved in post-translational modification of proteins with phosphate-containing compounds such as AMP (34).

In a previous study, insertions in MXAN_5793 suppressed the developmental defects caused by constitutive accumulation of Pxr-S (35), a processed non-coding small RNA (sRNA) that inhibits development when nutrients are present (36, 37). The mechanism of suppression is unknown, but MXAN_5793 has a predicted RNA-binding domain. Insertions in MXAN_5793 revealed previously unsuspected roles in both types of sporulation (Fig. 5A). Insertions in *rnd* (MXAN_5981) led to modest underrepresentation in both types of spore samples (Fig. S3A; Table S1). Rnd is RNase D, which is typically involved in 3' end processing of tRNA and other small, stable RNAs, and in *M. xanthus* is necessary for processing of the precursor sRNA that leads to Pxr-S (37). Several observations support a model in which Pxr-S, Rnd, and BsgA (the protease implicated in B-signal production) function in a nutrient-sensing pathway governing initiation of starvation-induced development (37). We speculate that Asg proteins also function in the pathway and that the pathway also controls glycerol-induced sporulation (Fig. S3B). In support, we note that insertions in *bsgA* (Fig. 2B), *asgA* (Fig. 5A), and *asgD* (MXAN_6996) and *spdR* (MXAN_1078) (Fig. S3A; Tables S1 and S2) positively and/or negatively affected fitness scores in both types of spore samples. AsgD is a hybrid RR-HK that appears to be involved in nutrient sensing (38). SpdR is an RR required for accumulation of the precursor sRNA that leads to Pxr-S (39). Insertions in *spdS* (MXAN_1077) encoding SpdR's cognate HK did not yield scores in our experiments. The processing that leads to Pxr-S may involve another RNase in addition to RNase D (37). Mining our data revealed two candidates, *rnc* (MXAN_3762) encoding ribonuclease III and *ybeY* (MXAN_4736) encoding an RNase typically involved in rRNA processing (Fig. S3A).

**Cell-envelope genes important for fruiting body sporulation**

Insertions in genes for several predicted membrane transporters and lipoproteins were depleted in starvation-induced spore samples (Fig. 6). MXAN_0441, 0983, and 3880 each have 12 predicted transmembrane regions (TMRs), but the molecules they transport across the IM are unknown. MXAN_0983 appears to be co-transcribed with upstream MXAN_0985/0984. Insertions in all three genes resulted in strong depletion in spore samples (Fig. 6). The proteins presumably form an envelope transporter with MXAN_0983 in the IM, MXAN_0984 in the periplasm, and MXAN_0985 in the outer membrane (OM). MXAN_3880 and upstream MXAN_3879 do not appear to be co-transcribed, but their insertions had similar fitness profiles (Fig. 6). MXAN_3879 is a predicted cytoplasmic orphan hybrid HK-RR-RR that may indirectly regulate MXAN_3880 transport activity during development. FadL is the OM porin for long-chain fatty acid uptake (40). A *fadL* mutant was proficient in development on starvation agar (41). In contrast, insertions in *fadL* (MXAN_7040) were enriched ~2-fold in late non-agg samples and depleted ~10-fold in time course 3 spore samples (Fig. 6), indicating a mild aggregation defect and a strong sporulation defect in submerged culture development of our Tn-Himar mutant pool. Insertions in MXAN_7039 had a similar fitness profile, possibly due to a polar effect, since this upstream gene appears to be co-transcribed with *fadL.* MXAN_7039 is a lipoprotein (40) broadly conserved in the Myxococcales, with a predicted lipase domain. MXAN_1162 is also a lipoprotein (40, 42) broadly conserved in the Myxococcales, with TPRs in a central LapB domain possibly involved in assembly of proteins for LPS biosynthesis, and C-terminal DUF3857 and YebA (transglutaminase or protease) domains. Insertions in MXAN_1162 were also depleted ~10-fold in time course 3 fruiting body spore samples (Fig. 6).

Insertions in *omrA* (MXAN_4426), MXAN_6629/6630/6631, and MXAN_1439 were slightly more enriched in agg samples than in non-agg samples, indicating enhanced aggregation, and these insertions were strongly depleted in starvation-induced spore samples (Fig. 6). OmrA has eight predicted TMRs and is postulated to flip aminoacylated phospholipids to the outer leaflet of the inner membrane, mediating a homeostatic response to OME (43) and conferring sensitivity to certain lipoprotein toxins (44). Developmental defects of an *omrA* mutant have not been reported previously. MXAN_6629 appears to be co-transcribed with downstream MXAN_6630/6631 and insertions in all three genes resulted in similar fitness profiles (Fig. 6). MXAN_6629 encodes a predicted UbiA family prenyltransferase with nine predicted TMRs and is likely involved in lipoquinone biosynthesis important for membrane electron transport/energy production *via* cellular respiration (45). MXAN_6630/6631 are predicted cytoplasmic oxidoreductases. MXAN_1439 appears to be co-transcribed with upstream MXAN_1438 for which insertions did not yield scores. MXAN_1439 has 11 predicted TMRs that form a putative glycosyltransferase RgtA/B/C/D-like domain.

Insertions in MXAN_5370 and MXAN_3746 had fitness profiles indicating mild aggregation defects and strong starvation-induced sporulation defects (Fig. 6). MXAN_5370 encodes a putative transmembrane glycosyltransferase, which together with downstream MXAN_5369 is likely co-transcribed from a weak promoter within *fadJ* (MXAN_5371) and a stronger promoter upstream of *fadI* (MXAN_5372). FadIJ are important for fatty acid oxidation, which generates energy from lipid bodies to produce spores with thicker coats and greater resistance to heat and UV light (46). However, a defined *fadI* insertion mutant in monoculture produced at least as many sonication-resistant spores as wild type (47). Insertions in *fadI* were not well represented in our pools and insertions in *fadJ* were not defective in spore formation (Table S2). Insertions in MXAN_3746 had a similar fitness profile as those in MXAN_5370 (Fig. 6). MXAN_3746 encodes a predicted cytoplasmic oxidoreductase and is likely co-transcribed with upstream MXAN_3748/3747, but insertions in these genes were not impacted in our experiments (Table S2). MXAN_3748 encodes serine palmitoyltransferase, which catalyzes the initial step of sphingolipid biosynthesis, and a defined MXAN_3748 insertion mutant in monoculture aggregated normally (48).

MXAN_1910 and downstream MXAN_1911 appear to be co-transcribed, and insertions in these genes were strongly depleted in starvation-induced spore samples and mildly depleted in glycerol-induced spore samples (Fig. 6). MXAN_1910 encodes a predicted LD-carboxypeptidase that may be important for the PG breakdown known to accompany both types of sporulation (33, 49). MXAN_1911 encodes a predicted serine hydrolase domain and is structurally similar to β-lactamases, perhaps providing protection against β-lactams during the vulnerable process of cellular shape change.

### Methods

#### Construction of a barcoded *M. xanthus* Himar transposon mutant library

We mutagenized *M. xanthus* DK1622 (50) by conjugation with *E. coli* donor APA752 harboring the pKMW3 barcoded himar transposon vector library (51). DK1622 from a freezer stock was streaked onto a CTT (1% Casitone, 10 mM Tris·HCl [pH 8.0], 1 mM $KH_2PO_4$-$K_2HPO_4$, 8 mM $MgSO_4$, [final pH 7.6]) 1.5% agar plate and incubated 4 days at 32°C. A yellow colony was inoculated into 5 mL CTTYE (CTT with 0.2% yeast extract) and the culture was shaken at 350 rpm for ~16 h at 32°C. The culture was diluted to 15-20 Klett units (KU) in 25 mL CTTYE and shaken as before until reaching 130-200 KU, the late-log phase of growth. In parallel, a 1-mL frozen stock of APA752 was thawed, inoculated into 20 mL Lysogeny Broth (LB) medium supplemented with 50 μg/mL kanamycin to select for the transposon-bearing plasmid and 0.3 mM diaminopimelic acid (DAP) to support growth of APA752, and shaken at 250 rpm for ~16 h at 37°C. The culture was diluted 50-fold into 200 mL LB Kan DAP and shaken as before until reaching 110 KU. The donor APA752 and recipient DK1622 cultures were centrifuged (7,000 × *g*, 5 min, 25°C), cells were suspended in LB DAP at 10,000 KU and 1,000 KU, respectively, by up-and-down pipetting and gentle vortexing, equal volumes were mixed (10:1 ratio of donor:recipient cells), and 0.5 mL aliquots were placed in the center of LB 1.5% agar DAP plates (90 mm diameter) and incubated at 32°C for 6 h to allow conjugation. The cell mixture from 8 plates was scraped into 5 mL CTT. Cells were suspended by up-and-down pipetting and gentle vortexing. To determine the mutation efficiency, a small portion of the cell suspension was diluted tenfold serially in CTT, and 0.2 mL aliquots of the undiluted suspension and the 10- and 100-fold dilutions were spread on CTT plates supplemented with 40 μg/mL kanamycin sulfate (Km40), while 0.2 mL aliquots of the $10^6$- and $10^7$-fold dilutions were spread on CTT plates to measure viable cells, with colonies counted after 4 days at 32°C. The mutation efficiency was ~2 × $10^{-5}$/viable *M. xanthus* cell. The remaining suspension was adjusted to 10% glycerol and stored at -80°C. The mutagenesis was repeated three times, for a total of four biological replicates. Based on the mutation efficiency, nine sub-libraries (three from one replicate and two from each of the others) estimated to contain ~25,000 mutants each, were generated by spreading 0.3 mL aliquots of thawed cell suspension from conjugation experiments on 10-13 CTT Km40 plates (150 mm diameter) per sub-library. After 4 days at 32°C, 2 mL CTTYE was added to each plate, cells were collected, suspended, pooled, adjusted to 10% glycerol, and stored in 1 mL aliquots at -80°C. One aliquot of each sub-library was thawed, the sub-libraries were pooled, the pool was diluted to 10 KU in CTTYE Km40 ug/ml kanamycin and the culture was shaken at 350 rpm and 32°C until reaching 120-140 KU. The culture was centrifuged (7,000 × *g*, 5 min, 25°C), cells were suspended in fresh CTTYE (20% of the original culture volume), adjusted to 10% glycerol, and stored in 1 mL aliquots at -80°C.

### Mapping Tn-Himar insertion sites in the *M. xanthus* mutant library

The mapping protocol we used has been described in detail (52). Briefly, genomic DNA was extracted, sheared to generate ~300-bp fragments, end-repaired, A-tailed, ligated to a custom Y adapter (Table S4), cleaned with SPRIselect beads using a two-sided (0.6×–0.25×) selection, and eluted from the beads. The Tn-Himar-containing fragments were enriched with a two-step nested PCR strategy using Q5 DNA polymerase with GC enhancer (New England Biolabs, M0491). The first reaction used the primers TS_pHimar_F+4 and TS_R with the following cycling parameters: 98°C for 3 minutes, 20× (98°C, 30 s; 66°C, 20 s; 72°C, 20 s), 72°C for 5 min, 4°C hold. The second reaction used the primes P5_TS F and P7_MOD_TS_index005 and the following cycling parameters: 98°C for 3 minutes, 15× (98°C, 30 s; 69°C, 20 s; 72°C, 20 s), 72°C for 5 min, 4°C hold (see Table S4 for primer sequences). The PCR products were cleaned with 1X SPRIselect beads (Beckman Coulter). The amplified fragments were sequenced at SeqCenter (Pittsburgh, PA) on one lane of a 10B 300 cycle NovaSeq X Plus flow cell (Illumina). The amplicons were loaded at a concentration of 100 pM and supplemented with approximately 40% phiX DNA.

Sequences were analyzed using custom scripts written and described by Wetmore and colleagues (51) and available at https://bitbucket.org/berkeleylab/feba/src/master/. Briefly, the locations of Tn-Himar insertions were aligned and mapped to the *M. xanthus* chromosome sequence (NCBI accession number CP000113.1) using BLAT, and unique barcode sequences were associated with their corresponding genome insertion location using the custom Perl script, MapTnSeq.pl. Sets of barcodes that reliably mapped to one location in the genome were identified using the custom Perl script, DesignRandomPool.pl with default parameters. Tn-Himar insertion mapping data are deposited at the NCBI Sequence Read Archive under BioProject accession PRJNA1517256.

#### Growth and development of the barcoded Tn-Himar mutant library

To grow the mutant library, a 1-mL frozen stock was thawed and inoculated into 10 mL CTTYE Km40 and shaken at 350 rpm for ~16 h at 32°C. The culture was diluted to 20 KU in 60 mL CTTYE Km40 and shaken as before until reaching 110 KU, the mid-log phase of growth. Fruiting body development in submerged culture with MC7 (10 mM morpholinepropanesulfonic acid [MOPS; pH 7.0], 1 mM $CaCl_2$) starvation buffer (53) was performed as described (54). Cells were harvested by centrifugation (7,000 × *g*, 5 min, 25°C) and the cell pellet was suspended in MC7 to 1,000 KU by up-and-down pipetting 15 times and vortexing 15 s, three times total. Input (0 h) samples (0.25 mL) of the 1,000-KU cell suspension were centrifuged (13,800 × *g*, 5 min, 25°C), supernatants were removed, and cell pellets were stored at -80°C. The 1000-KU cell suspension (0.25 mL) was added to MC7 (1.65 mL) in each well (35 mm diameter) of 6-well plastic plates. The plates were covered, wrapped with plastic wrap, and incubated at 32°C. Cells adhere to the bottom of the wells and undergo development. Samples were collected at indicated times by removing the supernatant from each well and adding 1 mL MC7, scraping cells off the well bottom with a 1-mL pipette tip, pelleting cells by centrifugation (13,800 × *g*, 5 min, 25°C), removing the supernatant and storing the cell pellet at -80°C. At the beginning of the third time course experiment, the mutant library starter culture was diluted to 20 KU in 70 mL CTTYE Km40 and shaken as before until reaching 80 KU, then 10 mL was removed for glycerol induction of sporulation, and the remaining culture was shaken as before until reaching 110 KU for fruiting body development. The 10-mL culture was adjusted to 0.5 M glycerol and shaken as before for 7 h, then samples (1.5 mL) were centrifuged (13,800 × *g*, 5 min, 25°C), supernatants were removed, and cell pellets were stored at -80°C. In all three experiments, for each sample, two biologically independent replicate time courses, each with three technical replicates, were collected.

**Separation of cells with different developmental fates**

Separation of non-agg from agg cells by low-speed centrifugation was performed as described (55). Briefly, cell pellets of samples collected as described above were suspended in 1 mL MC7 by up-and-down pipetting 15 times and vortexing 15 s, then centrifuged (50 × *g* for 5 min). The supernatant containing non-agg cells was carefully transferred to a different tube. The pellet containing agg cells remained. Both tubes were centrifuged (7,000 × *g*, 5 min, 25°C), supernatants were removed, and cell pellets were stored at -80°C. Spores were separated from non-spore agg cells by sonication to remove non-spore cells. Agg cell pellets from starvation-induced sporulation and cell pellets from glycerol-induced sporulation were suspended in 0.2 mL TE buffer (10 mM Tris·HCl pH 8.0, 1 mM EDTA) and 0.4 μL RNase A (10 mg/mL), then sonicated for 10 s three times with cooling on ice in between (using a Branson model 450 sonifier on setting 2 and a microtip, which was sufficient to lyse non-spore cells based on microscopic examination). To degrade non-spore DNA, 20 μL 10X reaction buffer (100 mM Tris·HCl pH 7.6, 25 mM $MgCl_2$, 5 mM $CaCl_2$) and 0.5 μL DNase I (10 mg/mL) was added and the mixture was incubated at 37°C for 10 min. To inactivate DNase I, 2 μL EDTA (0.5 M) was added and the mixture was incubated at 75°C for 10 min. To release spore DNA, 200 μL of the mixture was added to 200 mg acid-washed glass beads (100 μm diameter) and bead beating was performed for 2 min three times at maximum speed using a Mini-BeadBeater (Biospec). Spore DNA samples were stored at -20°C and used for PCR without further purification.

**Amplification and sequencing of Tn-Himar barcodes**

Genomic DNA was extracted from non-agg and agg cell pellets using guanidium thiocyanate as described (56). The DNA pellet was dissolved in 10-200 μL TE buffer and 1-4 μL were used as template (Table S5) in 20-μL barcode amplification reactions performed as described where the forward primer, Barseq_P1, was used for all templates and a uniquely indexed reverse primer, Barseq_P2_ITXXX, was used for each template (52) (see Table S4 for primer sequences). PCR products were separated on 2% agarose gels containing ethidium bromide to confirm amplification of the 190-bp product. Of note, using cells directly as described (52) did not amplify barcodes sufficiently for detection. We also note that guanidium thiocyanate extraction of spores (after DNase I digestion of non-spore DNA and inactivation of DNase I) did not lead to detectable barcode amplification (hence, the need for bead beating to release spore DNA as described above). Spore DNA samples (2 or 4 µL) were used as template (Table S5) in 20-µL barcode amplification reactions and PCR products were analyzed as above.

Based on visual approximation of product yield, reaction aliquots (4-10 µL) were pooled to roughly equalize the product amount from each reaction. The pool of PCR products was centrifuged and 90 µL of the mix was transferred to a fresh tube and cleaned with SPRIselect beads (Beckman Coulter) using a two-sided (1.2X–0.5X) selection as described (52). The amplified barcodes were sequenced at SeqCenter (Pittsburgh, PA) on one lane of a 10B 300 cycle NovaSeq X Plus flow cell (Illumina). The amplicons were loaded at a concentration of 100 pM and supplemented with approximately 40% phiX DNA. Barcode amplicon sequencing data for all experiments are available at the NCBI Sequence Read Archive under BioProject accession PRJNA1517256.

**Analysis of Tn-Himar strain fitness**

To calculate the fitness of the mutant strains in our experiments, we use the custom scripts generated by Wetmore and colleagues and described in (51). Briefly, the barcodes in each sample were counted and assembled using MultiCodes.pl and combineBarSeq.pl, then using the barcode abundance data and the mutant pool mapping information and the R script, FEBA.R, the fitness of each strain was calculated as a normalized $\log_2$ ratio of barcode counts in each developmental sample to counts in the 0 h sample of the same experiment. The fitness of each gene was calculated as the weighted average of strain fitness values, with the weight being inversely proportional to a variance metric (t-score) based on the total number of reads for each strain. Insertions in the first or last 10% of a gene were not included in gene fitness calculations. The complete set of gene-level individual fitness scores (f-scores) and t-scores for each developmental sample is listed in Table S3.

To identify genes that contribute to developmental phenotypes robustly, we focused on genes with absolute average t-scores >4 in any biological replicate sample from time courses 1, 2, or 3. This generated a list of 209 genes, a few of which had modest changes in fitness scores. We then eliminated genes for which the absolute average fitness score was <1 for all complete samples (i.e., 2 biological replicates, each with 3 technical replicates). This yielded a list of 200 genes (Table S1). These 200 genes were subjected to hierarchical clustering based on their average fitness scores. A heatmap of clustered fitness scores was rendered with Prism (GraphPad) with some manual rearrangement of genes to bring together genes that are adjacent on the chromosome and have similar fitness profiles. Subsets of this cluster, with additional manual rearrangements were generated to highlight key genes discussed in the text. Occasionally genes with similar patterns, but that did not meet the criteria for inclusion in Table S1, were manually added to the fitness heat maps presented in the text. Such genes are indicated with a # symbol.

**Tables in separate documents:**

**Table S1: Average fitness and t-scores for the set of genes that passed the selection criteria**

Excel file

**Table S2: Average fitness scores and t-scores for all genes**

Excel file

**Table S3: All scores for all genes from all samples**

Excel file

**Table S4 Primers and oligonucleotides**

| Primer / oligo | 5'-3' Sequence (description) | Notes |
|---|---|---|
| Y adapter oligos | | |
| Mod2_TruSeq | /5'P/GATCGGAAGAGCACACGTCTGAACTCCAGTCA<br>(5' phosphorylation) | Y-adapter (51) |
| Mod2_TS_Univ | ACGCTCTTCCGATC*T<br>(3' T is phosphorothioate bonded) | Y-adapter (51) |
| For two-step nested amplification of transposon junctions | | |
| TS_pHimar F+4 | ACACTCTTTCCCTACACGACGCTCTTCCGATCTNNNNNNCGCCCTGCAGGGATGTCCACGAGGTCT<br>(TruSeqRead1-hexamer-complementary to transposon U1 region) | Modified from Nspacer_barseq_pHimar (51) for nested amplification and to increase transposon-specific sequence |
| TS_R | GTGACTGGAGTTCAGACGTGTGCTCTTCCGATCT<br>(TruSeqRead2) | |
| P5-TS F | AATGATACGGCGACCACCGAGATCTACACTCTTTCCCTACACGACGCTCTTCCGATCT<br>(P5 sequence-TruSeqRead1) | |
| P7_MOD_TS_index5 | CAAGCAGAAGACGGCATACGAGAT**CACTGT**GTGACTGGAGTTCAGACGTGTGCTCTTCCGATCT<br>(P7 sequence-**index**-TruSeqRead2) | (51) |
| For amplification of barcodes | | |
| Barseq_P1 | AATGATACGGCGACCACCGAGATCTACACTCTTTCCCTACACGACGCTCTTCCGATCTNNNNNGTCGACCTGCAGCGTACG<br>(P5-TruSeqRead1-hexamer-complementary to transposon U2 region) | (51) |
| Barseq_P2_ITXXX | CAAGCAGAAGACGGCATACGAGATXXXXXXGTGACTGGAGTTCAGACGTGTGCTCTTCCGATCTGATGTCCACGAGGTCTCT<br>(P7 sequence-index-TruSeqRead2-complementary to transposon U1 region) | Uniquely indexed primers were used for each sample and are fully listed in (51) |

**Table S5 Barcode amplification samples**

| Time course 1 | | |
|---|---|---|
| Time point (h) | TE buffer (µL) | Template (µL) |
| 0 | 200 | 1 |
| 12 | 200 | 1 |
| 15 | 150 | 1 |
| 18 | 150 | 1 |
| 21 | 100 | 1 |
| 24 | 100 | 1 |
| 36 | 60 | 1 |

| Time course 2 | | | |
|---|---|---|---|
| Time point (h) | Sample | TE buffer (µL) | Template (µL) |
| 0 | | 100 | 1 |
| 12 | non-agg | 50 | 1 |
| | agg | 50 | 1 |
| 18 | non-agg | 20 | 1 |
| | agg | 50 | 1 |
| 24 | non-agg | 20 | 1 |
| | agg | 20 | 1 |
| 30 | non-agg | 20 | 2 |
| | agg | 20 | 2 |
| 36 | non-agg | 10 | 2 |
| | spore | NA[a] | 2 or 4 |
| 48 | non-agg | 10 | 2 |
| | spore | NA | 4 |
| 72 | non-agg | 10 | 4 |
| | spore | NA | 4 |

| Time course 3 | | | |
|---|---|---|---|
| Time point (h) | Sample | TE buffer (µL) | Template (µL) |
| 0 | | 100 | 4 |
| 7 | spore[b] | NA | 4 |
| 36 | agg | 10 | 4 |
| | spore | NA | 4 |
| 48 | agg | 10 | 4 |
| | spore | NA | 4 |
| 72 | agg | 10 | 4 |
| | spore | NA | 4 |

[a]Not applicable since spore DNA was not extracted using guanidium thiocyanate and therefore not precipitated and dissolved in TE buffer.
[b]Glycerol-induced spores

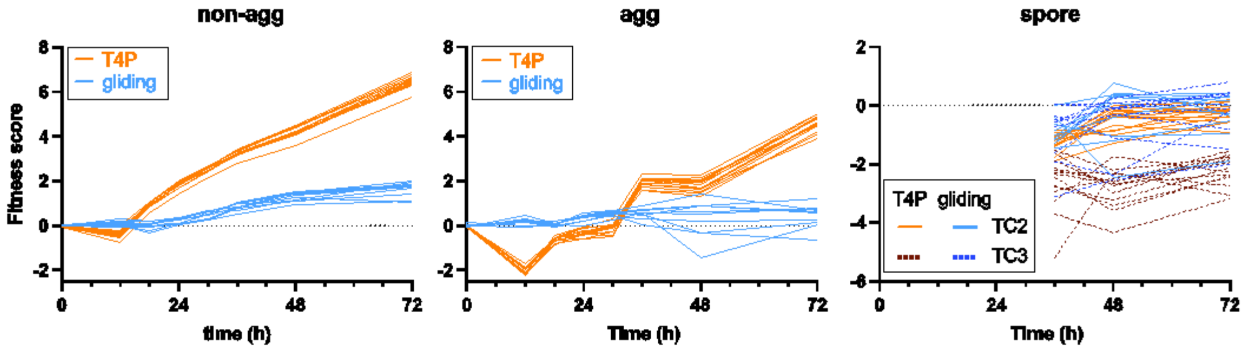


**FIG S1** T4P-dependent motility is more important than gliding motility for aggregation and sporulation. Comparison of fitness scores for genes involved in T4P-dependent motility (13 *pil* genes and *tgl*) and gliding motility (most *glt* genes, except *gltG*, and *cglB*) in non-agg, agg, and spore samples over time. Fitness scores of spore samples from both time course 2 (TC2) and time course 3 (TC3) are shown.

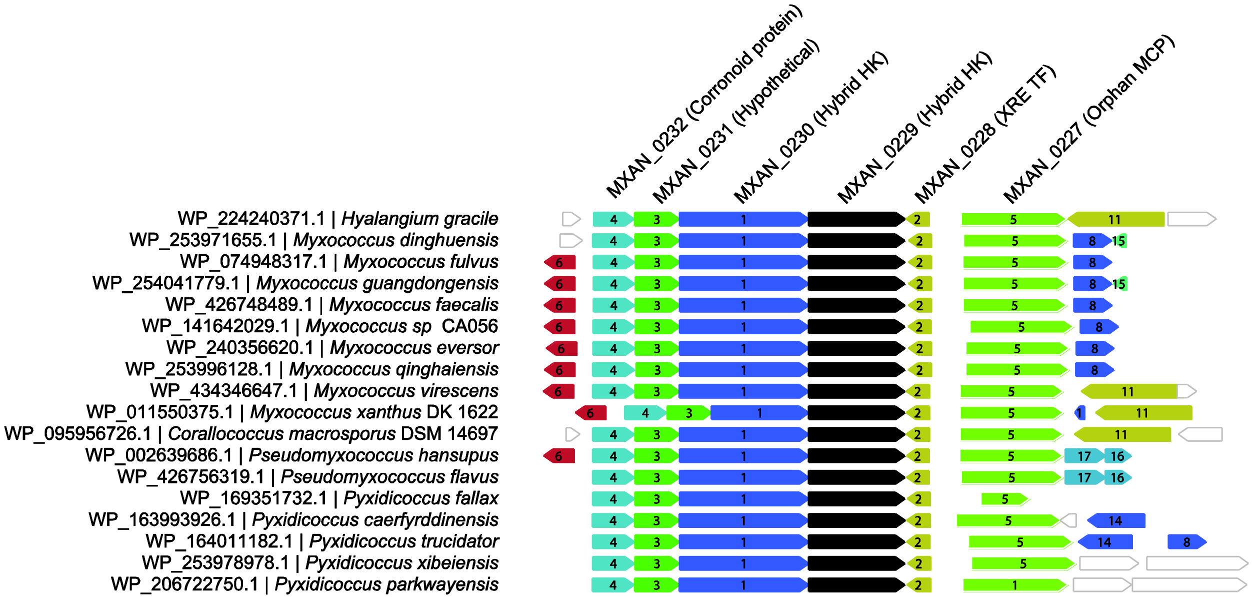


**FIG S2** Conserved gene neighborhood of MXAN_0229 homologs (webFlaGs/FlaGs2 output). Homologs of the *M. xanthus* MXAN_0229 protein were identified by BLASTP against the RefSeq Select database, and the genomic regions surrounding each hit were extracted and plotted as a gene-neighborhood map. Each row corresponds to one homolog of MXAN_0229 (RefSeq WP_ accession) and its source organism (left labels). The anchor gene (MXAN_0229 orthologue) is shown in black and aligned across rows. Neighboring ORFs are drawn as arrows (arrow direction indicates strand/orientation; arrow length reflects relative protein length). Colored/numbered arrows denote flanking-gene clusters inferred by FlaGs2 (same color/number = homologous flanking gene family across genomes), while uncolored/outlined arrows indicate genes not assigned to a conserved cluster in this set.

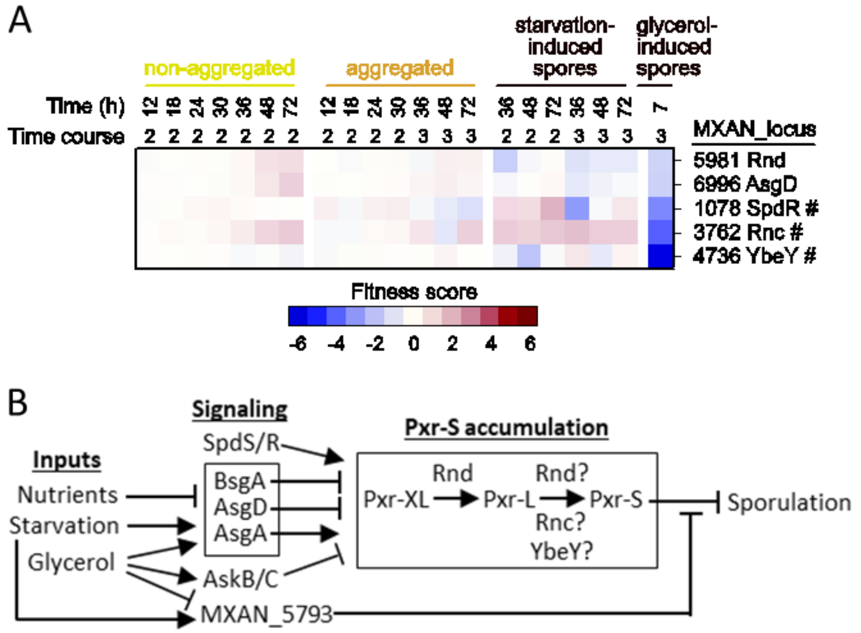


**FIG S3** Regulation of the small RNA Pxr-S that inhibits development when nutrients are present. (A) Heat map showing fitness scores of insertion mutants with known or inferred effects on Pxr-S function in sporulation. #; gene not among the 200 genes shown in Figure 1C and Table S1. (B) Model depicting the effects of inputs, signaling, and Pxr-S accumulation on sporulation. Inputs affect signaling protein activities that in turn regulate Pxr-S synthesis, processing, and/or stability (i.e., collectively Pxr-S accumulation), or in the case of MXAN_5793, inhibit Pxr-S without changing the Pxr-S level (35). Pxr-S inhibits sporulation. SpdR likely activates *pxr* transcription. Rnd processes Pxr-XL to Pxr-L. Rnd, Rnc, and YbeY are RNases that might process Pxr-L to Pxr-S. See the SI text and (37) for additional details and references.

### References


1. Yu R, Kaiser D. 2007. Gliding motility and polarized slime secretion. Mol Microbiol 63:454-467.
2. Pogue CB, Zhou T, Nan B. 2018. PlpA, a PilZ-like protein, regulates directed motility of the bacterium *Myxococcus xanthus*. Mol Microbiol 107:214-228.
3. Cho K, Treuner-Lange A, O'Connor KA, Zusman DR. 2000. Developmental aggregation of *Myxococcus xanthus* requires *frgA*, an *frz*-related gene. J Bacteriol 182:6614-6621.
4. Jakovljevic V, Leonardy S, Hoppert M, Sogaard-Andersen L. 2008. PilB and PilT are ATPases acting antagonistically in type IV pilus function in *Myxococcus xanthus*. J Bacteriol 190:2411-2421.
5. Bonner PJ, Black WP, Yang Z, Shimkets LJ. 2006. FibA and PilA act cooperatively during fruiting body formation of *Myxococcus xanthus*. Mol Microbiol 61:1283-1293.
6. Troselj V, Pathak DT, Wall D. 2020. Conditional requirement of SglT for type IV pili function and S-motility in *Myxococcus xanthus*. Microbiology (Reading) 166:349-358.
7. Bretl DJ, Muller S, Ladd KM, Atkinson SN, Kirby JR. 2016. Type IV-pili dependent motility is co-regulated by PilSR and PilS2R2 two-component systems via distinct pathways in *Myxococcus xanthus*. Mol Microbiol 102:37-53.
8. Wu SS, Kaiser D. 1997. Regulation of expression of the *pilA* gene in *Myxococcus xanthus*. J Bacteriol 179:7748-7758.
9. Caberoy NB, Welch RD, Jakobsen JS, Slater SC, Garza AG. 2003. Global mutational analysis of NtrC-like activators in *Myxococcus xanthus*: identifying activator mutants defective for motility and fruiting body development. J Bacteriol 185:6083-6094.
10. Li Y, Sun H, Ma X, Lu A, Lux R, Zusman D, Shi W. 2003. Extracellular polysaccharides mediate pilus retraction during social motility of *Myxococcus xanthus*. Proc Natl Acad Sci USA 100:5443-5448.
11. Willett JW, Tiwari N, Muller S, Hummels KR, Houtman JC, Fuentes EJ, Kirby JR. 2013. Specificity residues determine binding affinity for two-component signal transduction systems. mBio 4:e00420-13.
12. Ellehauge E, Norregaard-Madsen M, Sogaard-Andersen L. 1998. The FruA signal transduction protein provides a checkpoint for the temporal co-ordination of intercellular signals in *Myxococcus xanthus* development. Mol Microbiol 30:807-817.
13. Garza AG, Pollack JS, Harris BZ, Lee A, Keseler IM, Licking EF, Singer M. 1998. SdeK is required for early fruiting body development in *Myxococcus xanthus*. J Bacteriol 180:4628-4637.
14. Ogawa M, Fujitani S, Mao X, Inouye S, Komano T. 1996. FruA, a putative transcription factor essential for the development of *Myxococcus xanthus*. Mol Microbiol 22:757-767.
15. Pollack JS, Singer M. 2001. SdeK, a histidine kinase required for *Myxococcus xanthus* development. J Bacteriol 183:3589-3596.
16. Farrugia MA, Rajagopalan R, Kroos L. 2025. Transcriptomic analysis of *Myxococcus xanthus csgA, fruA, and mrpC* mutants reveals extensive and diverse roles of key regulators in the multicellular developmental process. BMC Genomics 26:355.
17. McLoon AL, Boeck ME, Bruckskotten M, Keyel AC, Sogaard-Andersen L. 2021. Transcriptomic analysis of the *Myxococcus xanthus* FruA regulon, and comparative developmental transcriptomic analysis of two fruiting body forming species, *Myxococcus xanthus* and *Myxococcus stipitatus*. BMC Genomics 22:784.
18. Stein EA, Cho K, Higgs PI, Zusman DR. 2006. Two Ser/Thr protein kinases essential for efficient aggregation and spore morphogenesis in *Myxococcus xanthus*. Mol Microbiol 60:1414-1431.
19. Cho K, Zusman DR. 1999. Sporulation timing in *Myxococcus xanthus* is contolled by the *espAB* locus. Mol Microbiol 34:714-725.

20. Higgs PI, Jagadeesan S, Mann P, Zusman DR. 2008. EspA, an orphan hybrid histidine protein kinase, regulates the timing of expression of key developmental proteins of *Myxococcus xanthus*. J Bacteriol 190:4416-4426.
21. Schramm A, Lee B, Higgs PI. 2012. Intra- and inter-protein phosphorylation between two hybrid histidine kinases controls *Myxococcus xanthus* developmental progression. J Biol Chem 287:25060–25072.
22. Shi X, Wegener-Feldbrugge S, Huntley S, Hamann N, Hedderich R, Sogaard-Andersen L. 2008. Bioinformatics and experimental analysis of proteins of two-component systems in *Myxococcus xanthus*. J Bacteriol 190:613-624.
23. Abellon-Ruiz J, Bernal-Bernal D, Abellan M, Fontes M, Padmanabhan S, Murillo FJ, Elias-Arnanz M. 2014. The CarD/CarG regulatory complex is required for the action of several members of the large set of *Myxococcus xanthus* extracytoplasmic function sigma factors. Environ Microbiol 16:2475-2490.
24. Bhagwat AS, Lieb M. 2002. Cooperation and competition in mismatch repair: very short-patch repair and methyl-directed mismatch repair in *Escherichia coli*. Mol Microbiol 44:1421-1428.
25. Kim JH, Kim H, Ko KS. 2026. Impact of DNA methyltransferases on bacterial fitness and genome stability in *Escherichia coli*. J Glob Antimicrob Resist 46:203-208.
26. Rolbetzki A, Ammon M, Jakovljevic V, Konovalova A, Sogaard-Andersen L. 2008. Regulated secretion of a protease activates intercellular signaling during fruiting body formation in *M. xanthus*. Dev Cell 15:627-634.
27. Velicer GJ, Kroos L, Lenski RE. 2000. Developmental cheating in the social bacterium *Myxococcus xanthus*. Nature 404:598-601.
28. Hoang Y, Franklin J, Dufour YS, Kroos L. 2024. Short-range C-signaling restricts cheating behavior during *Myxococcus xanthus* development. mBio 15:e0244024.
29. Schaal KA, Yu YN, Vasse M, Velicer GJ. 2022. Allopatric divergence of cooperators confers cheating resistance and limits effects of a defector mutation. BMC Ecol Evol 22:141.
30. Bernal-Bernal D, Abellon-Ruiz J, Iniesta AA, Pajares-Martinez E, Bastida-Martinez E, Fontes M, Padmanabhan S, Elias-Arnanz M. 2018. Multifactorial control of the expression of a CRISPR-Cas system by an extracytoplasmic function $\sigma$/anti-$\sigma$ pair and a global regulatory complex. Nucleic Acids Res 46:6726-6745.
31. Darnell CL, Wilson JM, Tiwari N, Fuentes EJ, Kirby JR. 2014. Chemosensory regulation of a HEAT-repeat protein couples aggregation and sporulation in *Myxococcus xanthus*. J Bacteriol 196:3160-3168.
32. Perez-Burgos M, Sogaard-Andersen L. 2020. Biosynthesis and function of cell-surface polysaccharides in the social bacterium *Myxococcus xanthus*. Biol Chem 401:1375-1387.
33. Ramirez Carbo CA, Irazoki O, Venkatesan S, Chen LJS, Morales HA, Garcia Avila AJ, Cheung HL, Cava F, Nan B. 2025. A novel mechanism for bacterial sporulation based on programmed peptidoglycan degradation. eLife 14:RP108250.
34. Roy CR, Cherfils J. 2015. Structure and function of Fic proteins. Nat Rev Microbiol 13:631-640.
35. Chen IK, Satinsky BM, Velicer GJ, Yu YN. 2019. sRNA-pathway genes regulating myxobacterial development exhibit clade-specific evolution. Evol Dev 21:82-95.
36. Yu YT, Yuan X, Velicer GJ. 2010. Adaptive evolution of an sRNA that controls *Myxococcus* development. Science 328:993.
37. Cossey SM, Velicer GJ, Yu YN. 2023. Ribonuclease D processes a small RNA regulator of multicellular development in myxobacteria. Genes 14:1061.
38. Cho K, Zusman DR. 1999. AsgD, a new two-component regulator required for A-signalling and nutrient sensing during early development of *Myxococcus xanthus*. Mol Microbiol 34:268-281.

39. Yu YN, Kleiner M, Velicer GJ. 2016. Spontaneous reversions of an evolutionary trait loss reveal regulators of a small RNA that controls multicellular development in myxobacteria. J Bacteriol 198:3142-3151.
40. Bhat S, Zhu X, Patel RP, Orlando R, Shimkets LJ. 2011. Identification and localization of *Myxococcus xanthus* porins and lipoproteins. PLoS One 6:e27475.
41. Bhat S, Boynton TO, Pham D, Shimkets LJ. 2014. Fatty acids from membrane lipids become incorporated into lipid bodies during *Myxococcus xanthus* differentiation. PLoS One 9:e99622.
42. Kahnt J, Aguiluz K, Koch J, Treuner-Lange A, Konovalova A, Huntley S, Hoppert M, Sogaard-Andersen L, Hedderich R. 2010. Profiling the outer membrane proteome during growth and development of the social bacterium *Myxococcus xanthus* by selective biotinylation and analyses of outer membrane vesicles. J Proteome Res 9:5197-5208.
43. Dey A, Wall D. 2014. A genetic screen in *Myxococcus xanthus* identifies mutants that uncouple outer membrane exchange from a downstream cellular response. J Bacteriol 196:4324-4332.
44. Vassallo CN, Sah GP, Weltzer ML, Wall D. 2021. Modular lipoprotein toxins transferred by outer membrane exchange target discrete cell entry pathways. mBio 12:e0238821.
45. Huang H, Levin EJ, Liu S, Bai Y, Lockless SW, Zhou M. 2014. Structure of a membrane-embedded prenyltransferase homologous to UBIAD1. PLoS Biol 12:e1001911.
46. Bullock HA, Shen H, Boynton TO, Shimkets LJ. 2018. Fatty acid oxidation is required for *Myxococcus xanthus* development. J Bacteriol 200:e00572-17.
47. Saha S, Kroos L. 2024. Regulation of late-acting operons by three transcription factors and a CRISPR-Cas component during *Myxococcus xanthus* development. Mol Microbiol 121:1002-1020.
48. Ring MW, Schwar G, Bode HB. 2009. Biosynthesis of 2-hydroxy and iso-even fatty acids is connected to sphingolipid formation in myxobacteria. Chembiochem 10:2003-2010.
49. Bui NK, Gray J, Schwarz H, Schumann P, Blanot D, Vollmer W. 2009. The peptidoglycan sacculus of *Myxococcus xanthus* has unusual structural features and is degraded during glycerol-induced myxospore development. J Bacteriol 191:494-505.
50. Kaiser D. 1979. Social gliding is correlated with the presence of pili in *Myxococcus xanthus*. Proc Natl Acad Sci USA 76:5952-5956.
51. Wetmore KM, Price MN, Waters RJ, Lamson JS, He J, Hoover CA, Blow MJ, Bristow J, Butland G, Arkin AP, Deutschbauer A. 2015. Rapid quantification of mutant fitness in diverse bacteria by sequencing randomly bar-coded transposons. mBio 6:e00306-15.
52. Fiebig A, Schnizlein MK, Pena-Rivera S, Trigodet F, Dubey AA, Hennessy MK, Basu A, Pott S, Dalal S, Rubin D, Sogin ML, Eren AM, Chang EB, Crosson S. 2024. Bile acid fitness determinants of a *Bacteroides fragilis* isolate from a human pouchitis patient. mBio 15:e0283023.
53. Kuner JM, Kaiser D. 1982. Fruiting body morphogenesis in submerged cultures of *Myxococcus xanthus*. J Bacteriol 151:458-461.
54. Rajagopalan R, Kroos L. 2014. Nutrient-regulated proteolysis of MrpC halts expression of genes important for commitment to sporulation during *Myxococcus xanthus* development. J Bacteriol 196:2736-2747.
55. Lee B, Holkenbrink C, Treuner-Lange A, Higgs PI. 2012. *Myxococcus xanthus* developmental cell fate production: heterogeneous accumulation of developmental regulatory proteins and reexamination of the role of MazF in developmental lysis. J Bacteriol 194:3058-3068.
56. Pitcher D, Saunders N, Owen R. 1989. Rapid extraction of bacterial genomic DNA with guanidium thiocyanate Lett Appl Microbiol 8:151–156.
